# RUNX2 inhibition disrupts a PAX3::FOXO1-RUNX2 feed-forward loop and dismantles oncogenic gene programs in fusion-positive rhabdomyosarcoma

**DOI:** 10.1101/2025.07.21.665972

**Authors:** Elizabeth A. Mendes, Aanandi Munshi, Archana Singh, Maisie D. Evans, Hsien-Chao Chou, Yong Yean Kim, Young Song, Ara Jo, Daniel Lee, Jaida Ciampi, Audrey Chambers, Samantha Weitzel, Michael Deel, Rex C. Bentley, Javed Khan, Darrell Green, Corinne M. Linardic

**Affiliations:** Division of Pediatric Hematology-Oncology, Department of Pediatrics, Duke University School of Medicine, Durham, NC, USA; Department of Pharmacology & Cancer Biology, Duke University School of Medicine, Durham, NC, USA; Norwich Medical School, University of East Anglia, Norwich Research Park, Norwich, United Kingdom; Amity Institute of Biotechnology, Amity Institute of Integrative Sciences and Health, Amity University Haryana, Gurugram, India; Oncogenomics Section, Genetics Branch, Center for Cancer Research, National Cancer Institute, NIH, Bethesda, MD, USA; Department of Biology, University of Evansville, Evansville, IN, USA; Department of Pathology, Duke University School of Medicine, Durham, NC, USA

**Author notes:** Target journal: Cancer Research (AACR). Corresponding author: Dr. Corinne M. Linardic, Division of Pediatric Hematology-Oncology, Department of Pediatrics, Duke University School of Medicine, Box 102382 DUMC, Durham, NC, 27710, USA.

**Keywords:** Rhabdomyosarcoma, RMS, PAX3::FOXO1, RUNX2, CADD522

## Abstract

Fusion-positive rhabdomyosarcoma is an aggressive pediatric cancer of skeletal muscle lineage, with a 5-year overall survival of <30% for high-risk disease, and <8% when metastatic. The *PAX3::FOXO1* fusion gene, resulting from *t*(2:13), is a signature driver of fusion-positive rhabdomyosarcoma, but similar to other transcription-factor based fusion genes in other cancers, not currently pharmacologically tractable. To identify novel druggable proteins in fusion-positive rhabdomyosarcoma tumor tissue and cell lines, we performed mRNA-seq of RMS patient tumors and utilizing the human FP-RMS cell lines Rh30 and Rh4, found that the RUNX2 transcription factor was the top druggable dependency. *In vitro* loss of function studies using genetic (RNAi) or pharmacologic (small molecule CADD522) inhibition showed that RUNX2 suppression inhibited FP-RMS cell growth, induced myogenic differentiation and apoptosis, and phenocopied PAX3::FOXO1 suppression. *In vivo* loss of function studies using conditional (dox-inducible) or pharmacologic (small molecule CADD522) blockade of tumor growth in a xenograft model system showed that RUNX2 suppression inhibited tumor growth. Mechanistically, we identify a PAX3::FOXO1 feed-forward loop whereby PAX3::FOXO1 binds a *RUNX2* enhancer to upregulate gene expression alongside MYOD1, while RUNX2 expression supports the expression of PAX3::FOXO1 at the mRNA and protein level.

**Significance:** RUNX2 inhibition reduces PAX3::FOXO1 expression and signaling, which impairs fusion-positive rhabdomyosarcoma oncogenic phenotypes. *In vivo* treatment with CADD522 decreased tumor growth and increased survival, indicating that RUNX2 is a promising therapeutic target.

## Introduction

Rhabdomyosarcoma (RMS) is the most common soft tissue sarcoma of childhood and adolescence, characterized by the expression of skeletal muscle markers (1). Current World Health Organization criteria subclassifies RMS by additional histopathologic features into four types: embryonal RMS (ERMS), alveolar RMS (ARMS), spindle cell/sclerosing RMS (SCRMS), and pleomorphic RMS (PRMS) (2). ERMS and ARMS are the most common, and now classified molecularly based on the absence (fusion-negative RMS; FN-RMS) or the presence (fusion-positive RMS; FP-RMS) of the PAX3/7::FOXO1 fusion oncogene, respectively. Despite these molecular advances, high risk RMS still portends a five-year overall survival of less than 30%, and metastatic RMS of less than 8% (3). Despite decades of research, the standard chemotherapy protocol for RMS remains vincristine, actinomycin D, and an alkylator such as cyclophosphamide or ifosfamide, drug combinations discovered in the 1970s. New therapies are desperately needed (4,5).

FP-RMS represents a disproportionally high percentage of high-risk RMS cases. The oncogenic driver of FP-RMS is PAX3::FOXO1, a chimeric transcription factor generated from the stable reciprocal translocation of chromosomes 2 and 13. PAX3::FOXO1 impairs myogenic differentiation via epigenetic dysregulation that produces myoblast-like cells capable of sustained proliferation but not terminal myogenic differentiation (6,7). Despite its role as an oncogenic driver, PAX3::FOXO1 is refractory to therapeutic targeting due to its lack of catalytic activity, and its intrinsically disordered structure, which leaves few pockets to bind small molecules (6,8,9). However, the transcriptional programs driven by PAX3::FOXO1 may yield alternate therapeutic targets. For example, prior work identified a PAX3::FOXO1 activated myogenic super-enhancer landscape that maintains the myoblastic state (8,10), including *MYCN* and *MYOD1* (8,10–12), and these are being evaluated as drug targets (13,14).

Here, we have taken a broad view of RMS to nominate additional members of this network that may be more amenable to clinical targeting. Using messenger (mRNA) sequencing data of primary patient tumor tissue, we compared the four histopathologic types of RMS, and with the DepMap resource, prioritized the transcription factors participating in this network. At the top of our list was RUNX2, a transcription factor previously identified as a PAX3::FOXO1 downstream target and part of the PAX3::FOXO1 interactome (8,10), but which has not been evaluated as a therapeutic target in FP-RMS.

RUNX2 is a member of the RUNX family (RUNX1, RUNX2, RUNX3) that is defined by a DNA-binding runt domain, and which largely operates as part of a heterodimeric core binding factor (CBF) complex with core binding factor-beta (CBFβ) to enhance DNA binding capacity (15,16). RUNX2 is best known for its essential role in the differentiation of mesenchymal stem cells (MSCs) into osteoblast and chondrocyte lineages. *RUNX2* transcription is regulated by two promoters, P1 and P2, with both isoforms expressed in osteoblast-lineage cells and chondrocytes, but expression is largely regulated by enhancers, which remain to be elucidated (15,16). RUNX2 is a master regulator and organizer, working in tandem alongside other transcription factors and co-factors (17). RUNX2 also has roles in the differentiation of bipotent MSCs into myotubes and osteogenic cells (18), but these are less well understood.

Here, we investigated the phenotypic consequences of RUNX2 inhibition in FP-RMS, the mechanism by which RUNX2 promotes downstream PAX3::FOXO1 signaling, and evaluated the consequences of RUNX2 inhibition *in vivo*. Using both genetic and pharmacological inhibition of RUNX2 *in vitro* and *in vivo*, we found that RUNX2 is essential to maintaining a proliferative FP-RMS cell state, while preventing apoptosis and terminal myogenic differentiation. Mechanistically, RUNX2 and PAX3::FOXO1 reciprocally regulate one another in a feed-forward loop to drive transcription of downstream PAX3::FOXO1 targets. Our findings suggest that directly inhibiting RUNX2 phenocopies PAX3::FOXO1 suppression and highlights its role as a legitimate druggable driver of oncogenic phenotypes in FP-RMS.

## Material and Methods

### Patient samples

The University of East Anglia Faculty of Medicine and Health Science Research Ethics Sub-Committee prospectively approved the collection and study of human RMS samples (ref: ETH2324-1212). Archived fresh frozen patient derived tissue samples (n=26) were obtained from the Royal Orthopaedic Hospital NHS Foundation Trust Research Tissue Bank (IRAS ID: 289182; REC ref: 22/EM/0042), which is a specialist tertiary hospital for sarcoma surgical management. Prior to tissue banking, a consultant histopathologist confirmed RMS histological interpretation using World Health Organization guidelines. This project included ARMS (n=5), ERMS (n=4), SCRMS (n=3), PRMS (n=1) and wildtype (control) skeletal muscle tissues (n=13). All individuals provided written informed consent to donate their tissue for research. Patient demographics are summarized in **Supplementary Table S1**.

### RNA-seq and bioinformatics of patient samples

RMS and matched adjacent control tissues were homogenized under liquid nitrogen. Total RNA was extracted using the miRNeasy Mini Kit (Qiagen, #217084). RNA samples were cleaned using the RNA Clean & Concentrator-5 kit (Zymo Research, #R1014). RNA concentration and integrity was measured on the Nanodrop 8000 spectrophotometer (Thermo Fisher Scientific, #ND-8000-GL) and the TapeStation (Agilent). RNA was stored at -80°C. mRNA libraries were constructed using the NEBNext ultra II RNA library prep kit (New England Biolabs, #E7775) and sequenced on a NovaSeq 6000 (Illumina) set to 150 bp paired end (PE) sequencing parameters. Small (sRNA) libraries were constructed using the NEBNext multiplex small RNA library prep kit (New England Biolabs) and sequenced on a NovaSeq 6000 (Illumina) set to 50 bp single end (SE) sequencing parameters.

For mRNA-derived cDNA libraries, FASTQ files were converted to FASTA files. Reads containing unassigned nucleotides were excluded. Trim Galore (RRID:SCR_011847) was used to remove adapter sequences and reads <20 nt. Trimmed reads were aligned to the human genome (v38) using HISAT2 (RRID:SCR_015530) (19). Transcripts were download from GENCODE (v46, RRID:SCR_014966) and Ensembl (v112, RRID:SCR_002344). Count matrices for transcripts were created using Kallisto (RRID:SCR_016583) (20). Differentially expressed genes were determined using the DESeq2 (v1.2.10, RRID:SCR_015687) package in RStudio (RRID:SCR_000432) (21–24). Differentially expressed genes were selected according to log2 fold change ≥2, p≤0.05 and false discovery rate (FDR) <5%. For sRNA derived cDNA libraries, FASTQ files were converted to FASTA. Reads containing unassigned nucleotides were excluded. Adapter sequences (3’: AGATCGGAAGAGCACACGTCT; 5’: GTTCAGAGTTCTACAGTCCGACGATC) were trimmed. sRNAs were mapped full length with 0 mismatches to the human genome (v38) and corresponding annotations using PatMaN (RRID_SCR_011831) (25). The latest set of human miRNAs were downloaded from miRBase (v22, RRID:SCR_003152) (26). Normalization and differential expression analysis was performed using DESeq2 (v1.2.10, RRID:SCR_015687) (22,23,27–32). Independent filtering was used to remove low abundance transcripts in normalized counts. sRNAs were considered differentially expressed if they had a p-value <0.05, FDR <5% according to the Benjamini–Hochberg procedure (22,25) and a log2 offset fold change >1.

### RNA-seq of FP-RMS cells

Total RNA was extracted from cells using RNeasy Plus Mini Kits (Qiagen, #74134). After QC, RNA-seq libraries were constructed using TruSeq Stranded mRNA kits (Illumina) and sequenced on a NextSeq2000 (Illumina) according to instructions of the manufacturer. Reads were mapped to the hg19 human reference genome using STAR (v.2.7.10a, RRID:SCR_004463), and gene expression was calculated as transcripts per million mapped reads (TPM) using RSEM (v.1.3.1, RRID:SCR_000262). For differential expression analysis, the DESeq2 (RRID:SCR_015687) package was utilized. Genes with adjusted *p*-value < 0.05 and fold change of 1.5 or above were considered significantly differentially expressed. GSEA analysis were performed on log2-fold change ranked list calculated by DESeq2’s (RRID:SCR_015687) lfcshrink function.

### Generation of cell lines and constructs

The human FP-RMS cell line Rh4 (RRID:CVCL_5916) were a gift from Dr. Beat Schaefer. Human FP-RMS cell lines Rh28 (RRID:CVCL_8725) and Rh30 (RRID:CVCL_0041) and human FN-RMS cell line RD (RRID:CVCL_1649), were gifts from Tim Triche (Children’s Hospital of Los Angeles, Los Angeles, CA). Rh4 (RRID:CVCL_5916), Rh30 (RRID:CVCL_0041), and Rh28 (RRID:CVCL_8725) cell lines express the *PAX3::FOXO1* fusion gene. U2OS (RRID:CVCL_0042) osteosarcoma cells were a gift from Dr. David Kirsch. SMS-CTR and RH36 human FN-RMS cell lines were gifts from Brett Hall (Columbus Children’s Hospital, OH, USA). All cell lines tested negative for mycoplasma (using Lonza MycoAlert PLUS test at the Duke University Cell Culture Facility) and were also authenticated by short tandem repeat analysis (Promega Powerplex 18D at Duke University DNA analysis facility) in 2021; Rh28 (RRID:CVCL_8725) and Rh30 (RRID:CVCL_0041) were reauthenticated in 2023. HEK293T (RRID:CVCL_0063) cells were obtained from the ATCC through the Duke University Cell Culture Facility and cultured as previously described (33). For all studies cells were imaged on a Leica DMi1 microscope and processed on Leica Application Suite Software. For visual clarity, image contrast was modified, and scale bars were added with ImageJ software (RRID:SCR_003070), quantification was not performed with adjusted images.

RNAi constructs to *RUNX2* (sh1, sh2, and sh4) were obtained from the Duke Functional Genomics Core Facility through their Sigma Mission TRC1 lentiviral shRNA Library. Knockdown (KD) constructs were stably expressed using established lentiviral and selection methods. Lentiviral particles were produced from HEK293T (RRID:CVCL_0063) cells transiently transfected with the lentiviral expression plasmid using FuGENE® 6 (Promega, #E2691). Lentiviral particles were harvested at 24 and 48 h post transfection and filtered through 0.45 µM filters. Polybrene was added to viral particles to a final concentration of 4 µg/mL. Rh4 and Rh30 cells were selected with 1.0 µg/mL puromycin (Sigma) for 72 h. RNAi constructs to negative control (QIAGEN, #1022076) and *PAX3::FOXO1* (Qiagen, #1027423) were purchased and transfected with lipofectamine RNAiMAX (Thermo Fisher Scientific, #13778150) according to manufacturer recommendations and collected after 48 h. RNAi (shRNA and siRNA) sequence information is listed in **Supplementary Table S2A,B.**

Doxycycline inducible constructs were cloned using the same oligo sequences from the TRC1 lentiviral shRNA library. Forward and reverse oligonucleotides were ordered with overhangs and loop sequences from Integrate DNA Technologies (IDT) and were then annealed together by PCR according to manufacturer recommendations. The tet-pLKO-puro (Addgene, #21915, RRID:Addgene_21915) backbone was cut using restriction enzymes AgeI and EcoRI (New England Biolabs). The vector and the insert (annealed oligos) were then ligated (TaKaRa, #6023) and transformed using Stbl3 cells. Colonies were picked and grown in lysogeny broth (LB) with ampicillin. Plasmids were extracted (Zymo, #D4213), sequenced by Eton Biosciences, and aligned with SnapGene (RRID:SCR_015052) to ensure that constructs were cloned correctly. Since the AgeI cut site is destroyed in the cloning process, additional verification of cloning included a restriction enzyme digestion with EcoRI, AgeI, and BamHI.

### Quantitative real time PCR

RNA was isolated from cell pellets stored at -80°C using a RNeasy Mini Kit (Qiagen, #74106) according to manufacturer specifications. Conversion to cDNA was completed using the Omniscript RT kit (Qiagen, #205113). Quantitative real-time PCR (qRT-PCR) was performed using the SYBR Green system (Bio-Rad) according to manufacturer specifications. Primer sequence information used for qRT-PCR are listed in **Supplementary Table S2C.** Each biological experiment was run in triplicate. Technical and/or biological replicates are specified in figure legends.

### Pharmacologic agents

Computer aided drug design molecule 522 (CADD522) (SelleckChem, #S0790) was solubilized in DMSO to a 10µM stock. Further dilutions were made in PBS.

### ApoTox-Glo triplex assay

Viability and apoptosis assays were performed using the ApoTox-Glo Triplex Assay (Promega, #G6320) according to manufacturer specifications. Rh30 cells were plated at 5 x 10^3^ cells/well in a 96 well plate in triplicate, samples were read 72 h following the completion of puromycin selection (72 h) with the SpectraMax i3x Multi-Mode Microplate Reader to measure the transmittance and fluorescence.

### Immunoblotting

Cell pellets were lysed with high detergent RIPA (Cell Signaling Technology, #9806S) and Halt™ protease and phosphatase inhibitor cocktail (Thermo Fisher Scientific, #78442) then passaged through a 21-gauge needle for optimal shearing of DNA or sonicated for 3-5 sec using the Branson Fisher Scientific 150E Sonic Dismembrator at 22.5 kHz. Protein concentration was measured using the protein assay dye reagent concentrate (Bio-Rad, #500006). 4X protein Sample loading buffer (Thermo Fisher Scientific, #LC2570 and #NP0007) was combined with 20% β-mercaptoethanol or sample reducing agent (Thermo Fisher Scientific, #B0009) and then added to protein samples and then lysates were boiled for five min at 95°C. 20-30 µg of lysate was loaded onto 10% or 4%-15% gradient Mini-PROTEAN precast gel (Bio-Rad) and resolved with SDS-PAGE and then transferred to Immobulin-FL polyvinylidene difluoride membrane (Millipore, #IPFL00010) or nitrocellulose membrane (Bio-Rad, #1620112). Membranes were immunoblotted with primary monoclonal antibodies anti-RUNX2 (Cell Signaling Technology, #8486S, RRID:AB_10949892), anti-FOXO1 (Cell Signaling, #2880S, RRID:AB_2106495) and anti-β-ACTIN (Cell Signaling Technology, #4970, RRID:AB_2223172). Polyclonal antibodies used included anti-GAPDH (Thermo Fisher Scientific, #PA1-9046, RRID:AB_1074703). Membranes were reacted with a secondary antibody, either anti-rabbit or anti-goat and then fluorescence measured on the Odyssey (Li-Cor) or imaged by enhanced chemiluminescence on the Chemi-Doc (Bio-Rad). Depending upon protein abundance, West Pico PLUS (Thermo Fisher Scientific, #34580) or West Atto Ultimate Sensitivity (Thermo Fisher Scientific, #A38556) substrates were used and imaged at various exposures according to manufacturer protocol. Uncropped immunoblots are provided for review purposes in **Supplemental Presentation SP1.**

### Colony formation

750 cells/well (Rh30) were plated in a 6 well plate and incubated at 37°C and 5% CO_2_ for 20 days. Media was changed every other day. After the incubation period, cells were stained in a 0.1% (w/v) crystal violet solution with 25% MeOH to fix the cells for one 1 h. Cells were then washed with water for 10 min twice and left to dry overnight. The stained colonies were quantified using the GelCount machine (Oxford Optronix).

### Cell cycle analysis

Cells were collected 72 h after selection and fixed with ice cold 70% ethanol. Cells were centrifuged and the ethanol was removed. Cells were resuspended in 500 µL of PBS and propidium iodide and RNase A was mixed into solution slowly to avoid cell lysis. Cell cycle analysis was preformed using the Canto Flow Cytometry machine through the Duke Flow Cytometry Core. Data was analyzed using FlowJo (RRID:SCR_008520).

### MF20 staining

Myogenic differentiation was assessed by MF20 staining under light microscopy, as previously described (33). In brief, cells were plated at equal density to ∼60% confluence in 6-well plates in regular media (RPMI with 10% fetal bovine serum). Once adherent, cells were treated with CADD522 for 72 h. Myotubes were fixed, permeabilized, and stained with primary antibody anti–sarcomere-myosin hybridoma MF20 followed by secondary biotinylated anti-mouse IgG, then HRP-streptavidin with a 3,3′-diaminobenzidine reagent. MF20 was obtained from the Developmental Studies Hybridoma Bank, created by the NICHD of the NIH and maintained at The University of Iowa, Department of Biology, Iowa City, IA 52242.

### Growth curves

Cell growth was assayed using Trypan blue staining followed by automated cell counting on the TC20™ automated cell counter (BioRad). Cells were cultured in 6 well dishes and counted at three time points between days 1 through 3 in duplicate. 2 × 10^5^ cells were plated per replicate at each time point.

### *In vivo* xenograft assays

Xenograft studies utilized 8 × 10^6^ (doxycycline-induced *RUNX2* KD experiment) or 6 × 10^6^ (CADD522 experiment) cells resuspended in growth factor replete Matrigel (BD Biosciences, #354234) and implanted subcutaneously into the flanks of immunodeficient SCID/*beige* mice. Once tumors were palpable (doxycycline-induced *RUNX2* KD experiment) or 200 mm^3^ (CADD522 experiment), mice were randomly divided into two groups followed by vehicle or compound administration. For doxycycline studies, mice received regular chow (Purina) or doxycycline containing chow (Enivgo). Prior to use, the CADD522 compound was dissolved in 5%DMSO/95%PBS at 10 mg/kg and injected intraperitoneally (IP). Mice in the vehicle group received 5%DMSO/95%PBS solution via IP injection. Tumors were measured 3 times weekly using calipers and tumor volume calculated as [((width × length)/2)^3^]/2. For doxycycline studies mice were sacrificed at specified timepoints of 13 days (sh2) and 15 days (sh4), upon reaching an Institutional Animal Care and Use Committee (IACUC)-defined maximum tumor burden (2,000 mm^3^) or decline in health. In the CADD522 studies, mice were sacrificed upon reaching an Institutional Animal Care and Use Committee (IACUC)-defined maximum tumor burden which was established as tumors measuring >15 mm in one direction, >12 mm in two directions, or decline in health rather than at a single timepoint for survival analyses. Portions of tumors were preserved in formalin-fixed and paraffin-embedded (FFPE) for immunohistochemistry or flash frozen in liquid nitrogen for RNA extraction using RNeasy Mini Kit (Qiagen, #74134).

### Hematoxylin and eosin

For all xenografts hematoxylin and eosin (H&E) staining, paraffin-embedded sections were deparaffinized in xylene, rehydrated through graded ethanols, and then submerged into hematoxylin (Cancer Diagnostics, #16600). Next, samples were submerged in water, clearing solution (Cancer Diagnostics, #16703-C), water, bluing solution (Cancer Diagnostics, #16702), water, eosin (Cancer Diagnostics, #16601), dehydrated through graded ethanols, and xylene using the Leica Autostainer XL.

### Immunohistochemistry

CADD522 xenografts were stained with rabbit monoclonal Ki-67 (Cell Signaling Technology, #12202, RRID:AB_2620142) according to the manufacturer’s protocol. Briefly, paraffin-embedded sections were deparaffinized in xylene, rehydrated through graded ethanols, and then submerged into citrate unmasking solution for heat-induced antigenic retrieval, blocked with animal-free blocking solution (Cell Signaling Technology, #15019), incubated with Ki-67 (Cell Signaling Technology, #12202, RRID:AB_2620142) primary antibody (1:800) 4 °C overnight and developed using the Signalstain® Boost IHC Dec. (HRP, Rab) (Cell Signaling Technology, #8114S) and DAB (Signalstain® DAB diluent, Cell Signaling Technology, #11724S; Signalstain® DAB chromogen concentrate, Cell Signaling Technology, #11725S) followed by counterstaining with Mayer’s hematoxylin (Sigma Aldrich, #MSH16-500), dehydration, clearing and mounting. Sections were captured on a Leica DMLB microscope with DFC425 camera and processed on Leica Application Suite software. For quantification, 4 random fields at 40X magnification per section were captured and analyzed by manual counting with the aid of ImageJ (RRID:SCR_003070) software. For visual clarity, image contrast was modified, and scale bars were added with ImageJ software (RRID:SCR_003070), quantification was not performed with adjusted images.

Doxycycline-inducible xenografts were stained with rabbit monoclonal Ki-67 (Abcam, #AB16667, clone SP6, RRID:AB_302459). The antibody is listed as for research use only. Ki-67 (Abcam, #AB16667, clone SP6, RRID:AB_302459) was used at 1:200 dilution with Ab Diluent (Discovery, #760-108). Immunohistochemistry tests were performed using the Ultra Discovery automated staining platform. The tissue sections were pretreated (epitope retrieval) with cell conditioning solution CC1 (Roche, #950-124) for 56 min, then incubated with Ki-67 (Abcam, #AB16667, clone SP6, RRID:AB_302459) for 1 h at 36 °C. Rabbit IgG, substituted for the primary antibody, was used as the negative control. After binding of the primary antibody, anti-rabbit HQ (Roche, #760-4815, RRID:AB_2811171) were applied and incubated for 12 min, followed by 12 min incubation with anti-HQ HRP (Roche, #760-4820, RRID:AB_3068525) for antigen detection. Roche HRP Detection is a biotin-free, hapten (HQ) anti-hapten (anti-HQ) linked horseradish peroxidase-linker antibody conjugate system for the detection of tissue-bound primary antibodies. The kit includes peroxidase blocking reagent, post primary IgG linker reagent (HQ labelled secondary antibody), and anti-HQ-HRP reagent to localize HQ labelled secondary antibodies, the immunohistochemistry reactions were visualized with DAB chromogen and counterstained hematoxylin. Sections were photographed under an optical microscope, 4 random fields per section were used to analyze with ImageJ (RRID:SCR_003070) software.

For all xenografts, Cleaved Caspase 3 (CC3) staining, the rabbit monoclonal CC3 antibody (Cell Signaling Technology, #9661S, RRID:AB_2341188) was used at 1:800, with Ab Diluent (Discovery, #760-108). Immunohistochemistry tests were performed using the Ultra Discovery automated staining platform. The tissue sections were pretreated (epitope retrieval) with cell conditioning solution CC1 (Roche, #950-124) for 56 min. CC3 (Cell Signaling Technology, #9661S, RRID:AB_2341188) was applied and incubated for 60 min at 36°C. Rabbit IgG, substituted for the primary antibody, was used as the negative control. After binding of the primary antibody, anti-rabbit HQ (Roche, #760-4815) were applied and incubated for 12 min, followed by 12 min incubation with anti-HQ HRP. For visualization, ChromoMap DAB (Discovery, #760-159) was applied and incubated for 5 min. Roche HRP Detection is a biotin-free, hapten (HQ) anti-hapten (anti-HQ) linked horseradish peroxidase-linker antibody conjugate system for the detection of tissue-bound primary antibodies. The kit includes peroxidase blocking reagent, post primary IgG linker reagent (HQ labelled secondary antibody), and anti-HQ-HRP reagent to localize HQ labelled secondary antibodies. The immunohistochemistry reactions were counterstained with hematoxylin. Sections were imaged on a Leica DMLB microscope with DFC425 camera and processed on Leica Application Suite software, 4 random fields per section were used to analyze with ImageJ (RRID:SCR_003070) software.

### Statistical analysis

All statistical analysis, except for sequencing, were performed using GraphPad Prism (RRID:SCR_002798) software. A *p-*value of 0.05 or lower was considered significant, and *p*-values were considered significant at *, *p*< 0.05; **, *p*<0.01; ***, *p*< 0.001; and ****, *p*<0.0001. Unless otherwise notes, data is presented as the mean and standard error (SE). Statistical tests used include unpaired student’s t-test with Welch’s correction when appropriate (cell-based assays), Pearson’s correlation coefficient when appropriate, two-way analysis of variance (ANOVA) or mixed model with the Holm-Šídák correction (cell growth curves), one-way ANOVA with the Dunnett correction (cell-based assays), multiple unpaired student’s t-test with a two-stage linear set-up procedure of Benjamini, Krieger, and Yekutieli methods applied (CADD522 *in vivo* FDR<0.1, doxycycline-inducible *in vivo* FDR<0.01), survival analysis evaluated by Gehan-Breslow-Wilcoxon test. For sequencing experiments of patient samples, variability between sequencing libraries was evaluated using scatter plots, replicate-to-replicate differential expression size split box plots, intersection and Jaccard similarity analysis. Empirical differential expression analysis was confirmed by parametric (t-tests) and non-parametric (Mann-Whitney-U) tests. All sequencing data presented in this study fulfilled log2 fold change >1, *p*<0.05 and FDR ≤5% criteria. For CRISPR DepMap data, the box plot of the CERES dependency score, which is based on DepMap data from cell depletion assays in CRISPR screens, demonstrating a specific dependency for *RUNX2* in fusion-positive ARMS compared to all cancer types analyzed together. A lower CERES score indicates a higher likelihood that the gene of interest is essential in a given cell line. A score of 0 is equivalent to a gene that is not essential whereas a score of -1 corresponds to the median of all common essential genes. Two-group comparisons were performed in parallel across genes using the LIMMA (RRID:SCR_010943) RStudio (RRID:SCR_000432) package (34), which uses parametric empirical Bayes methods to pool information across genes when assessing the significance of observed group differences. *P*-values for each gene are computed from empirical Bayes moderated t-statistics. For sequencing experiments of FP-RMS cell lines, genes with adjusted *p*-value < 0.05 and fold change of 1.5 or above were considered significantly differentially expressed.

## Data availability

All data supporting the findings of this study are available within the article and Supplementary files or from the corresponding author on request. Raw sequencing files are available at Gene Expression Omnibus (GEO) (RRID:SCR_005012) under the accessions GSE279954 (mRNA) and GSE280045 (sRNA). Raw sequencing files for Rh30 and Rh4 *RUNX2* KD are available at GSE298855 (mRNA).

## Results

### *RUNX2* is highly expressed in RMS primary tumor tissue

To investigate clinically relevant dysregulated genes in RMS, establish an additional primary human tumor dataset to aid RMS researchers, and to identify additional targetable proteins important for the FP-RMS myogenic circuit, we analyzed 13 flash-frozen patient-derived RMS primary tumor samples, including alveolar (ARMS), embryonal (ERMS), spindle cell/sclerosing (SCRMS), and pleomorphic (PRMS) histologies, combined with 13 patient-derived non-malignant wildtype skeletal muscle samples (SkM) (**Supplementary Table S1**) using mRNA-seq. Around 29M reads per sample were obtained (**Supplementary Table S1**). Short reads were mapped to the human genome, which revealed that the majority mapped to protein coding exons (**Fig. 1A**). Pearson’s correlation coefficient (PCC) showed a linear correlation between the RMS and SkM samples (**Fig. 1B**), meaning there was a difference in mRNA expression patterns between tumor and normal tissues. The coefficient of determination of the SkM group was high (R2<0.8), confirming tissue similarity. RMS intragroup variation was not strongly correlated, demonstrating a high variation in mRNA expression patterns between tumor samples (**Fig. 1B**). There was a low correlation between the PRMS sample and all other samples including SkM and RMS, with the strongest correlation with the SCRMS sample (R2=0.588) (**Fig. 1B**). Processed sequencing data was reduced to 2-dimensions, where we observed distinct clustering across the samples (**Fig. 1C**). Most RMS subtype groups overlapped, indicating a similarity in differentially expressed genes; however, PRMS was distinctly separate from all other samples suggesting that PRMS is transcriptionally different from other RMS subtypes (**Fig. 1C**), though this was a single patient sample.

**Figure 1.**
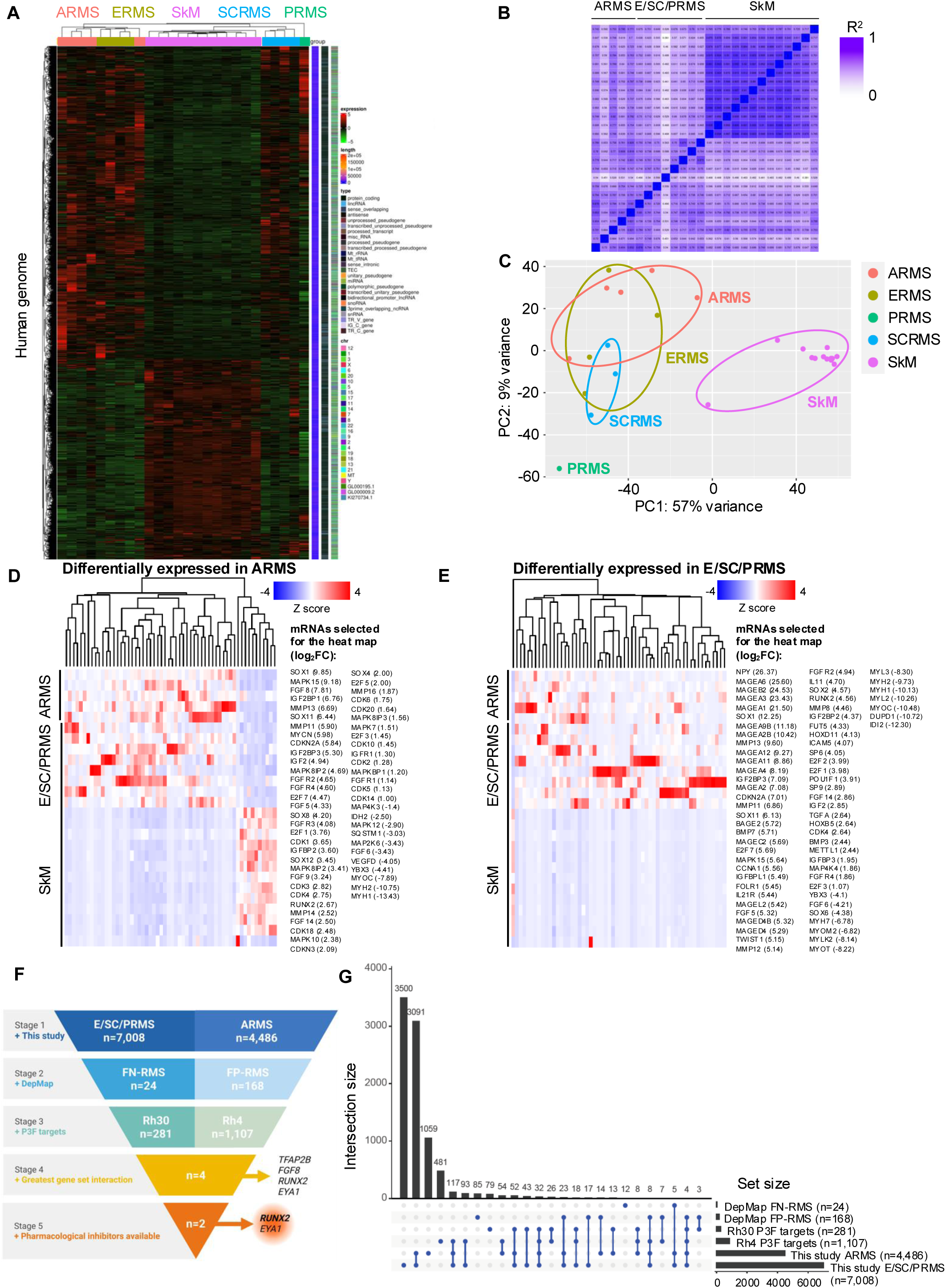
mRNA-seq performed on archival fresh frozen RMS tissues classified according to their histological interpretation. **A,** Quality check of mapped short read sequencing reads to the human genome in n=5 ARMS (sample IDs: R01, R02, R05, R08, R13) n=4 ERMS (sample IDs: R07, R09, R10, R12), n=3 SCRMS (sample IDs: R06, R14, R14), n=1 PRMS (sample ID: R11) and n=14 SkM (sample IDs: R17-28 and R30) tissues. Most of the reads mapped to protein coding mRNA loci. **B,** Pearson correlation coefficient (PCC) plot shows a linear correlation between the sample sets. **C,** Data reduction to two-dimensions via biplot principal component analysis (PCA) shows consistent groups along the PC1 axis that correspond to ARMS (light red), ERMS (olive), SCRMS (blue), PRMS (green) and SkM (pink) tissues. **D,E** Heat map based hierarchical cluster analysis of selected differentially expressed genes (x-axis) across ARMS and combined ERMS, SCRMS, and PRMS tissues (y-axis) when compared to the SkM with the log_2_ fold change in expression also shown. Z-score refers to high (red) and low (blue) gene expression using normalized values when compared to the mean of total sequencing reads. **F,** Candidate selection of differentially expressed genes across the current study (stage 1), the cancer DepMap (stage 2), Rh30 and Rh4 P3F targets (stage 3). Candidate genes of interest included those with the greatest gene set intersection (stage 4) and with pharmacological inhibitors (stage 5) available. Top candidate is bolded. **G,** UpSet plot to visualize intersected genes (and therefore generate gene/s-of-interest for downstream studies) across the current study, the cancer DepMap, and Rh30 and Rh4 cell line P3F targets.

To identify and visualize differentially expressed genes in ARMS samples, which are typically fusion-positive, heat map-based hierarchical cluster analysis was performed (**Fig. 1D**). Processed data for all genes is provided in **Supplementary Table S3**. Differentially expressed genes for the heat map were included according to their relevance to sarcoma/cancer (8,35). Among the selected differentially expressed genes, 47 were upregulated and 11 were downregulated (**Fig. 1D**). The most abundant upregulated genes in ARMS samples were *SOX1* (log_2_ fold change (FC)=9.86), *MAPK15* (log_2_ FC=9.18) and *FGF8* (log_2_ FC=7.82) (**Fig. 1D**) FP-RMS related genes (8,10–12) of interest that were also upregulated included oncogenes such as *MYCN*, *IGF2*, *RUNX2*, *MMP13* and *MMP11* (**Fig. 1D**). The most markedly downregulated were *MYH1* (log_2_ FC=-13.43), *MYH2* (log_2_ FC=-10.77) and *MYOC* (log_2_ FC=-7.9) (**Fig. 1D**).

Differentially expressed genes in ERMS, SCRMS and PRMS, which are typically (but not always) fusion-negative were also visualized through heat map-based hierarchical cluster analysis (**Fig. 1E**). Processed data for all genes is provided in **Supplementary Table S3**. The most abundant upregulated genes included *NPY* (log_2_ FC=26.38), *MAGEA6* (log_2_ FC=25.6) and *MAGEB2* (log_2_ FC=24.53) (**Fig. 1E**). The most markedly downregulated genes included *IDI2* (log_2_ FC=-12.31), *DUPD1* (log_2_ FC=-10.72) and *MYOC* (log_2_ FC=-10.48) (**Fig.1E**). The *MAGE* gene family, which is being evaluated as a candidate for immunotherapy (36), was the most represented gene group with 14 differentially expressed genes (**Fig. 1E**).

Next, all ARMS and E/SC/PRMS differentially expressed genes were compared to a series of independent publicly available datasets to identify potential FP-RMS and FN-RMS therapeutic and mechanistic targets of interest (**Fig. 1F**). Gene set intersection size of processed data was determined by comparing this study’s differentially expressed genes against RMS cell line dependencies and known Rh30 and Rh4 high confidence PAX3::FOXO1 targets, identified through CUT&RUN and ChIP-seq experiments following PAX3::FOXO1 degradation or genetic inhibition (**Fig. 1F,G**) (8,10). Out of 11,494 differentially expressed genes identified in ARMS (n=4,484) and E/SC/PRMS (n=7,008), 27% or 3,091 genes were exclusively represented in both datasets (**Fig. 1G**). When the data was narrowed further to account for potential fusion-status, 0 differentially expressed E/SC/PRMS genes were exclusive to FN-RMS and not FP-RMS dependencies, but 5 genes, *IGF2BP1*, *OSTC*, *CEP152*, *CNPY2*, and *LMO4*, were exclusive to FN-RMS dependencies as well as ARMS and E/SC/PRMS differentially expressed genes (**Fig. 1G; Supplementary Tables S3A, S4A**). These hits were not pursued further because we were primarily interested in the gene candidates that were exclusive to probable fusion-status in addition to RMS histological interpretation. This latter analysis identified a subset of 4 differentially expressed genes in ARMS and E/SC/PRMS, *TFAP2B*, *FGF8*, *EYA1*, and *RUNX2*, exclusive to 4/5 gene sets, the FP-RMS DepMap, Rh30 FP-RMS targets, and Rh4 FP-RMS targets (**Fig. 1F,G; Supplementary Tables S3A, S4A**). To further narrow this list, we cross-referenced with the literature to determine whether pharmacological inhibitors were either available or in development. We found that *EYA1* (log_2_ FC ARMS=2.23 E/SC/PRMS=2.06) and *RUNX2* (log_2_ FC ARMS=2.67 E/SC/PRMS=4.56) fit these criteria (**Fig. 1F**) (31,37–39). There are several EYA1 inhibitors in development including DS-1-38, which blocks sonic hedgehog signaling, but despite the incidence of EYA1 in Ewing’s sarcoma there were no publications using these inhibitors in sarcomas (6,40). Alternatively, the RUNX2 inhibitor, computer aided design molecule 522 (CADD522) reduces the DNA-binding capacity of RUNX2 and has been tested *in vitro* and *in vivo* in bone sarcoma and breast cancer models (31,38,39,41). Due to the increased differential expression of *RUNX2* versus *EYA1,* we moved forward to study RUNX2 as a therapeutic target in FP-RMS.

## *RUNX2* dependency is specific to pediatric sarcomas including RMS

To evaluate the clinical relevance of RUNX2 and its validity as a potential therapeutic and mechanistic RMS target of interest, we interrogated publicly available patient databases. Using the St. Jude PeCan database (https://pecan.stjude.cloud) that contains a cohort of 2,486 pediatric tumors, we found *RUNX2* was highly expressed in FP-RMS (**Supp. Fig. S1A**), second only to expression in the bone sarcoma osteosarcoma (OS), where the role of RUNX2 is well characterized (42). *RUNX2* expression was highest in FP-RMS, followed by FN-RMS, while non-sarcoma pediatric cancers including neuroblastoma (NBL), glioma (GLM), ependymoma (EPDY) and medulloblastoma (MBL) were less dependent on *RUNX2* (**Supp. Fig. S1A**). mRNA-seq cluster analysis was used to visualize *RUNX2* expression specifically in sarcomas, where the highest expression was in FP-RMS and osteosarcoma (**Supp. Fig. S1B**). Further analysis of a published dataset showed that *RUNX2* expression was higher in FP-RMS tumors when compared to FN-RMS and myoblasts (**Supp. Fig. S1C**) (8). Analysis of a separate, non-overlapping dataset from the Oncogenomics website (https://omicsoncogenomics.ccr.cancer.gov/cgi-bin/JK) hosted by the NCI showed a significant association of poor survival with high expression of *RUNX2* in all RMS regardless of fusion status and histology (*p*=0.0143) (**Supp. Fig. S1D)** and combined FP-RMS and FN-RMS cases with known fusion status (*p*=0.0399) (**Supp. Fig. S1E)**. When separated and analyzed by fusion presence, the statistical power was weaker, but trending towards significance, possibly due to the lower number of patients in the high *RUNX2* group, FP-RMS (*p*=0.0554) high *RUNX2* (n=10) and low *RUNX2* (n=40) (**Supp. Fig. S1F**). In the case of FN-RMS (*p=*0.1421), despite a higher number of patients high *RUNX2* (n=19) and low *RUNX2* (n=45) groups (**Supp. Fig. S1G**), statistical power was further from significance than in FP-RMS. Thus, we conclude that high *RUNX2* expression was correlated with poor survival across all RMS cases and may portend an even worse outcome in *PAX3/7::FOXO1* expressing patients (**Supp. Fig. S1E-G**). More data is required to establish a strong link between high *RUNX2* expression, fusion status, and survival.

Single-nucleus RNA sequencing (snRNA-seq) of GTExPortal data of normal skeletal muscle tissue showed that 2.43% of skeletal muscle myocyte cells express *RUNX2* (**Supp. Fig. S2A**). To gain insight into the dependency of individual RMS cell lines on *RUNX*2, we interrogated the DepMap CRISPR database and found that *RUNX2* is strongly selective in human FP-RMS cell lines, ranking #51 in FP-RMS alone (**Supplementary Table S5A**) and #77 compared to all other cancer cell lines (**Supp. Fig S2B; Supplementary Table S5B**) (43). Human FP-RMS cell lines exhibit a significantly higher dependency on *RUNX2* compared to FN-RMS and other solid tumor types (**Supp. Fig. S2C**) (43). Immunoblot analyses validated elevated RUNX2 protein levels in FP-RMS cell lines compared to FN-RMS and positive control OS cell lines (**Supp. Fig. S2D**). In this work, due to their dependency on and high expression of RUNX2 (**Supp. Fig. S2C,D**), the majority of experiments going forward were performed in Rh30 cells, with supporting complementary experiments performed in Rh4 cells.

Last, since RUNX1 is a close ortholog of RUNX2, in parallel we investigated the expression and dependency of RUNX1 in RMS according to publicly available datasets. When comparing FP-RMS to FN-RMS tumors and normal myoblasts, *RUNX1* expression was lower (**Supp. Fig. S3A,B**) which was consistent with increased *RUNX1* expression in E/SC/PRMS patients (log_2_ FC=1.44), but not in ARMS (**Fig. 1D**) (6). Though *RUNX1* expression was elevated in FP-RMS and FN-RMS cell lines (**Fig. S3A**) and ARMS and ERMS patients (**Supp. Fig. S3B**), Kaplan-Meier analyses revealed that increased *RUNX1* expression was associated with increased survival in all RMS (*p*=0.0640) (**Supp. Fig. S3C**) (10).

### *In vitro* KD of *RUNX2* in FP-RMS cell lines impairs oncogenic phenotypes

Combining the information gained from analysis of RMS patient samples, the role of RUNX2 in other human cancers, and publicly available patient and cell line-based RMS datasets, we hypothesized that by interfering with RUNX2 activity we could impair FP-RMS oncogenic phenotypes and potentially disrupt PAX3::FOXO1 signaling. To achieve RUNX2 loss of function, we generated four lentiviral constructs bearing unique short hairpin (sh)RNAs to the *RUNX2* coding sequence. Because sh3 was ineffective in suppressing *RUNX2* levels, it was excluded from further analysis. Stable expression of any of the three remaining shRNAs (sh1, sh2, or sh4) resulted in RUNX2 suppression as assessed by qRT-PCR and immunoblot in Rh30 and Rh4 cells (**Fig. 2A,B; Supp. Fig. S4A,B**). Phenotypically, *RUNX2* KD significantly reduced cell growth over the course of 72 h after selection in Rh30 and Rh4 cells (**Fig. 2C; Supp. Fig. S5C**). *RUNX2* KD also impaired colony formation of Rh30 cells (**Fig. 2D,E**). To gain insight into Rh30 and Rh4 mechanism of decreased cell growth, we imaged FP-RMS cells after RUNX2 suppression and found that RUNX2 KD resulted in decreased cell confluency accompanied by changes in cell morphology (**Fig. 2F; Supp. Fig. S5D**). Specifically, cells appeared elongated and myotube-like or had cellular blebbing and fragmentation consistent with cell death. Parallel studies in Rh4 cells replicated these morphologies (**Fig. 2F; Supp. Fig. S5D**). To verify the differentiation phenotype visualized in cell culture, we performed qRT-PCR analysis, which revealed that *MYOD1* and *OC,* genes associated with differentiation of MSCs, myoblasts, and osteoblasts, were significantly upregulated following *RUNX2* KD in Rh30 (**Fig. 2G,H**) (44).

**Figure 2.**
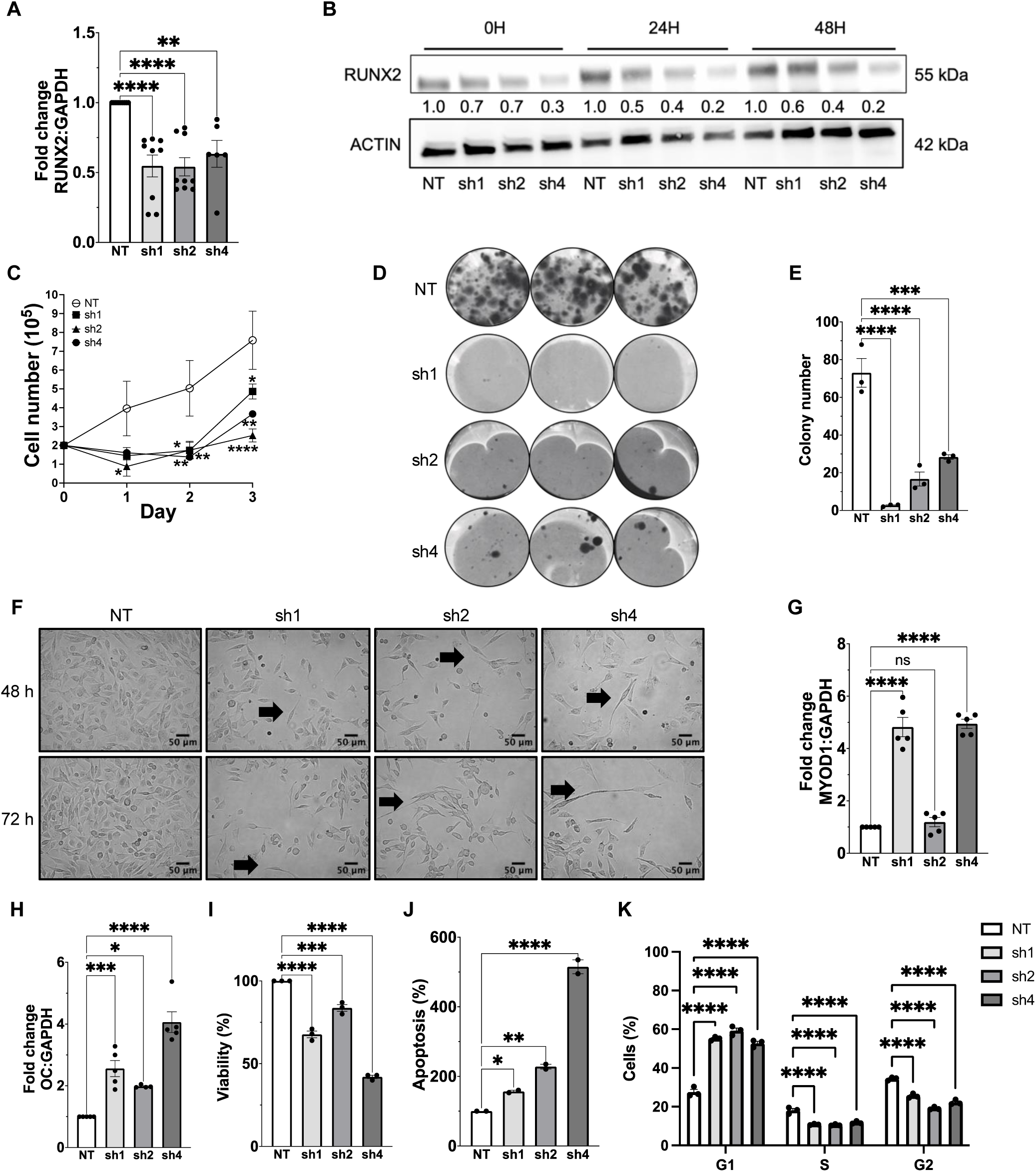
*RUNX2* knockdown in human FP-RMS cells (Rh30) impairs classical oncogenic phenotypes *in vitro*. **A,** qRT-PCR for *RUNX2* quantification in the negative control (NT) Rh30 cells and the three independent knockdown cells (sh1, sh2, sh4). Each black dot represents an independent biological replicate. **B,** Normalized RUNX2 protein expression in the negative control (NT) Rh30 cells and the three independent knockdown cells (sh1, sh2, sh4) measured at 0 h, 24 h and 48 h. **C,** Cell growth in negative control and *RUNX2* stable knockdown Rh30 cells. Black dots are independent biological replicates. **D,** Colony formation in negative control and *RUNX2* stable knockdown Rh30 cells. **E,** Colony numbers in negative control and *RUNX2* stable knockdown Rh30 cells. Black dots are independent technical replicates. **F,** *RUNX2* knockdown through three independent stably expressed shRUNX2 lentiviral preparations (sh1, sh2, sh4) plus negative control (shNT) in Rh30 cells. Black arrows point to visible areas of differentiation. Cells were imaged at 20X magnification and captured at 48 h and 72 h. Scale bars are 25 µm. **G,H,** qRT-PCR for known drivers of mesenchymal stem cell differentiation; *MYOD1* and *OC*, following *RUNX2* stable knockdown. Black dots are independent biological replicates. **I,** Cell viability of negative control and *RUNX2* stable knockdown Rh30 cells. Black dots are independent technical replicates. **J,** Quantification of caspase 3/7 mediated apoptosis in negative control and *RUNX2* stable knockdown Rh30 cells. Black dots are independent technical replicates. **K,** Cell cycle analysis in negative control and *RUNX2* stable knockdown Rh30 cells. Black dots are independent biological replicates consisting of three technical replicates each.

Increased *MYOD1* expression was not significant with sh2 (FC=1.89) while upregulation of *OC* was less robust in sh2 (FC=1.98) versus sh1 (FC=2.56) and sh4 (FC=4.06) (**Fig. 2G,H**). To verify the apoptotic phenotype visualized in cell culture, we performed cell viability assays paired with apoptosis assays (**Fig. 2I,J**). sh4 demonstrated a >50% decrease in cell viability and a 4-fold increase in apoptosis compared to the negative control (NT) (**Fig. 2I,J**). To further evaluate whether a reduction in proliferation, in addition to increased apoptosis, was responsible for decreased viability, we performed cell cycle analysis of Rh30 cells (**Fig. 2K**). Across all shRNA constructs, we saw an increased accumulation of approximately 40% of cells in G1 compared to the negative control (**Fig. 2K**). Also, there was a significant decrease of cell in S phase by approximately 50% and G2/M by approximately 10% across all shRNAs (**Fig. 2K**). Together, these data support the importance of *RUNX2* expression in maintaining oncogenic phenotypes required for FP-RMS cell growth, cell state, and cell survival.

### Bulk transcriptome analysis of *RUNX2* KD validates phenotypic findings and suggests impaired PAX3::FOXO1 activity

To identify the global genetic changes responsible for oncogenic phenotypes following *RUNX2* KD in Rh30 and Rh4 cells, we performed mRNA-seq of FP-RMS cells with stable sh4 RUNX KD at three timepoints: 0 h, 24 h, and 48 h after completion of selection (**Fig. 3**). This analysis not only aimed to identify changes in gene expression patterns, but also evaluated how expression fluctuates over time, to inform future RUNX2 mechanistic and pharmacologic studies. Around 20M reads per sample were obtained (**Supplementary Table S6**). Short reads were mapped to the human genome, which confirmed the majority mapped to protein coding exons (**Supplementary Tables S6, S7**). Processed sequencing data was reduced to 2-dimensions where we observed distinct clustering across the samples (**Fig. 3A,B**). PC1 accounted for 54% of variance while PC2 accounted for an additional 27% variance in Rh30 (**Fig. 3A**). PC1 accounted for 37% of variance while PC2 accounted for an additional 23% variance in Rh4 (**Fig. 3B**). Even though Rh30 and Rh4 are two distinct FP-RMS cell lines, the majority of *RUNX2* KD groups overlapped at distinct duplicate timepoints, indicating a similarity in differentially expressed genes; however, 0 h post selection was distinctly separate from 24 h and 48 h suggesting that downstream effects of *RUNX2* KD amplify over time (**Fig. 3A,B**).

**Figure 3.**
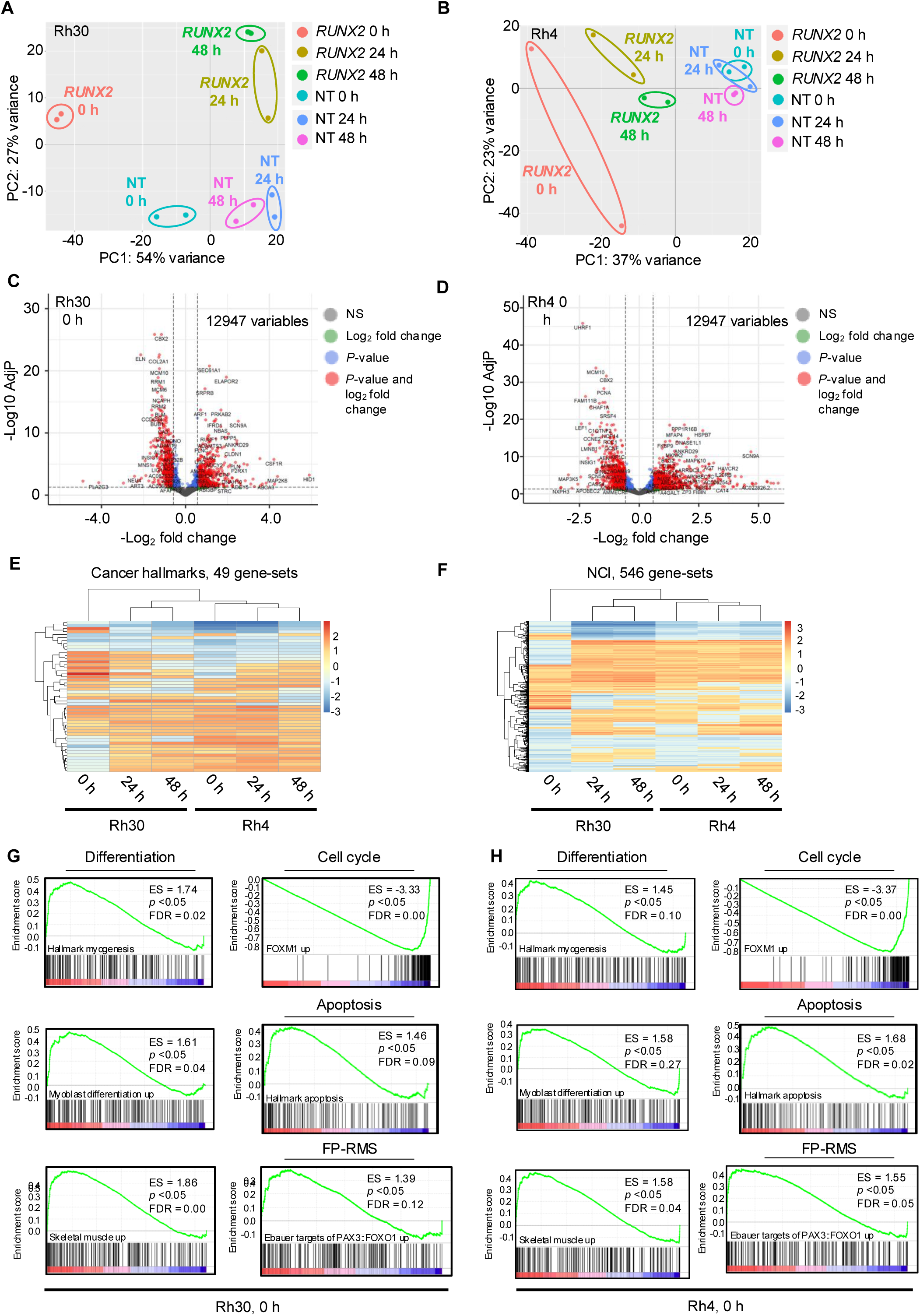
Transcriptomic alterations following *RUNX2* stable knockdown *in* vitro. **A,B,** Data reduction to two-dimensions via biplot PCA shows consistent groups along the PC1 axis that correspond to Rh30 and Rh4 cells *RUNX2* knockdown at 0 h post selection (light red), 24 h post selection (olive), 48 h post selection (green), and negative control (NT) cells at identical time points (turquoise-0 h, blue-24 h, and pink-48 h). **C,D,** Differentially expressed genes between shNT and sh4 *RUNX2* knockdown Rh30 and Rh4 cells at 0 h post selection. X-axis= -Log_2_FC. Y-axis= -Log10 AdjP. **E,F,** Heat map based hierarchical cluster analysis following gene set enrichment analysis (GSEA) of Rh30 and Rh4 cells at selected timepoints (x-axis), and cancer hallmark (n=49) and NCI (n=546) gene-sets (y-axis). Key represents normalized enrichment score (NES). **G,H** Enrichment plots for Rh30 and Rh4 cells following *RUNX2* knockdown at 0 h post selection. Selected gene-sets characterize the effect of *RUNX2* knockdown on differentiation, cell cycle, apoptosis, and FP-RMS regulatory circuitry. Enrichment score (ES), *P*-values, and false discovery rate (FDR) are shown.

Of the differentially expressed genes in Rh30 (**Fig. 3C**), the most abundant upregulated genes following *RUNX2* KD at 0 h, 24 h, and 48 h post selection include *HID1* (log_2_ FC=5.90), *COL6A6* (log_2_ FC=3.80) and *C4BPB* (log_2_ FC=6.09) respectively (**Supplementary Table S7**). Genes with known roles in FP-RMS biology were also downregulated at all time points including *MYCN*, *FGFR4*, *NOTCH1*, *WWTR1,* and *RUNX2* (**Supplementary Table S7**) (1,12,45). Of the differentially expressed genes in Rh4 (**Fig. 3D**), the most abundant upregulated genes following *RUNX2* KD at 0 h, 24 h, and 48 h post selection include *KLHDC7B* (log_2_ FC=5.82), *ACP5* (log_2_ FC=4.35) and *SCN9A* (log_2_ FC=4.15) respectively (**Supplementary Table S7**). Genes of interest with known roles in promoting FP-RMS that were also downregulated at all time points included the same genes as in Rh30 except for *MYCN* (**Supplementary Table S7**).

Differentially expressed genes in *RUNX2* KD samples were visualized via volcano plot (**Fig. 3C,D**). Differentially expressed genes were included in a Gene Set Enrichment Analysis (GSEA) to confirm the key roles of RUNX2 in FP-RMS. Two selected cancer hallmark and NCI gene-sets captured at 0 h, 24 h, and 48 h after selection largely became more enriched overtime in Rh30 and Rh4 (**Fig. 3E,F**). Processed data for additional GSEA analysis is provided in **Supplementary Table S8**. Gene sets involved in regulating myogenesis, skeletal muscle differentiation, myoblast differentiation, and apoptosis had positive enrichment scores that strengthened overtime following *RUNX2* KD in Rh30 and Rh4 (**Fig. 3G,H**). Alternatively, gene-sets involved in positively regulating cell cycle, including FOXM1, had negative scores (**Fig. 3G,H**) (46). Studies have reported that FOXM1 is a promising RMS therapeutic target (47), and these results suggest that *RUNX2* inhibition secondarily inhibits a FOXM1 gene-set signature. The most interesting enrichment plot conveyed that *RUNX2* KD upregulated the same gene-set signature that is upregulated with *PAX3::FOXO1* KD, suggesting that RUNX2 inhibition may genetically and morphologically phenocopy direct inhibition of the FP-RMS driver (**Fig. 3G,H**). These findings highlight the importance of *RUNX2* and its impact on FP-RMS biology.

### Conditional genetic *RUNX2* KD decreases *PAX3::FOXO1* expression *in vitro* and impairs tumor growth *in vivo*

To prepare for evaluation of RUNX2 KD *in vivo*, we engineered conditional (dox-inducible) shRNAs to RUNX2. Hairpin sequences sh2 and sh4 targeting *RUNX2* were subcloned into a doxycycline-inducible backbone and validated *in vitro* with Rh30 FP-RMS cells (**Fig. 4A,B**). Cells were cultured and treated with a range of doxycycline doses for 72 h; doxycycline-inducible KD phenocopied constitutively active KD (**Fig. 4C**; **Fig. 2**). Since mRNA-seq of *RUNX2* KD lead to increased gene expression of targets increased with *PAX3::FOXO1* KD, we performed qRT-PCR for *PAX3::FOXO1* and found *RUNX2* inhibition downregulated expression of the fusion transcript (**Fig. 4D**). To determine the consequences of *RUNX2* inhibition *in vivo*, cells were grown and injected subcutaneously into the flanks of SCID/*beige* mice (**Fig. 4E; Supp. Fig. S5A,B**). Mice with *RUNX2* KD had significantly reduced tumor volume compared to control mice, suggesting that RUNX2 inhibition attenuates oncogenic phenotypes *in vivo* (**Fig. 4E; Supp. Fig. S5A,B**). Tumors harvested from mice with sh2 *RUNX2* KD weighed significantly less than control tumors, while those from mice with sh4 RUNX2 did not meet statistical significance, likely because tumors began to escape the growth suppressive KD effects (**Fig.4E; Supp. Fig. S5A,B**). H&E of sh2 tumors did not show histological differences (**Fig. 4G, top row**). IHC for Ki-67 and CC3 was performed to confirm whether the anti-proliferative and pro-apoptotic effects seen *in vitro* hold true *in vivo;* proliferation was not significantly decreased, but one tumor exhibited less Ki-67 staining. The effect on apoptosis was significantly increased by approximately 5-fold in all sh2 tumors with active *RUNX2* KD compared to control tumors (**Fig. 4G-I**). qRT-PCR for select genes evaluated *in vitro* were repeated with RNA from sh2 tumors (**Fig. 4J**). *RUNX2* and *PAX3::FOXO1* expression decreased by ∼40% for a subset of tumors (**Fig. 4J**) and *MYOD1* and *OC* expression increased by 4-fold in some instances (**Fig. 4J**), the variability in response is likely due to compensation of *RUNX2* transcription towards conclusion of the experiment. Across all primers and all tumors, *OC* expression was significantly increased validating that some genetic changes observed *in vitro* are replicated *in vivo*.

**Figure 4.**
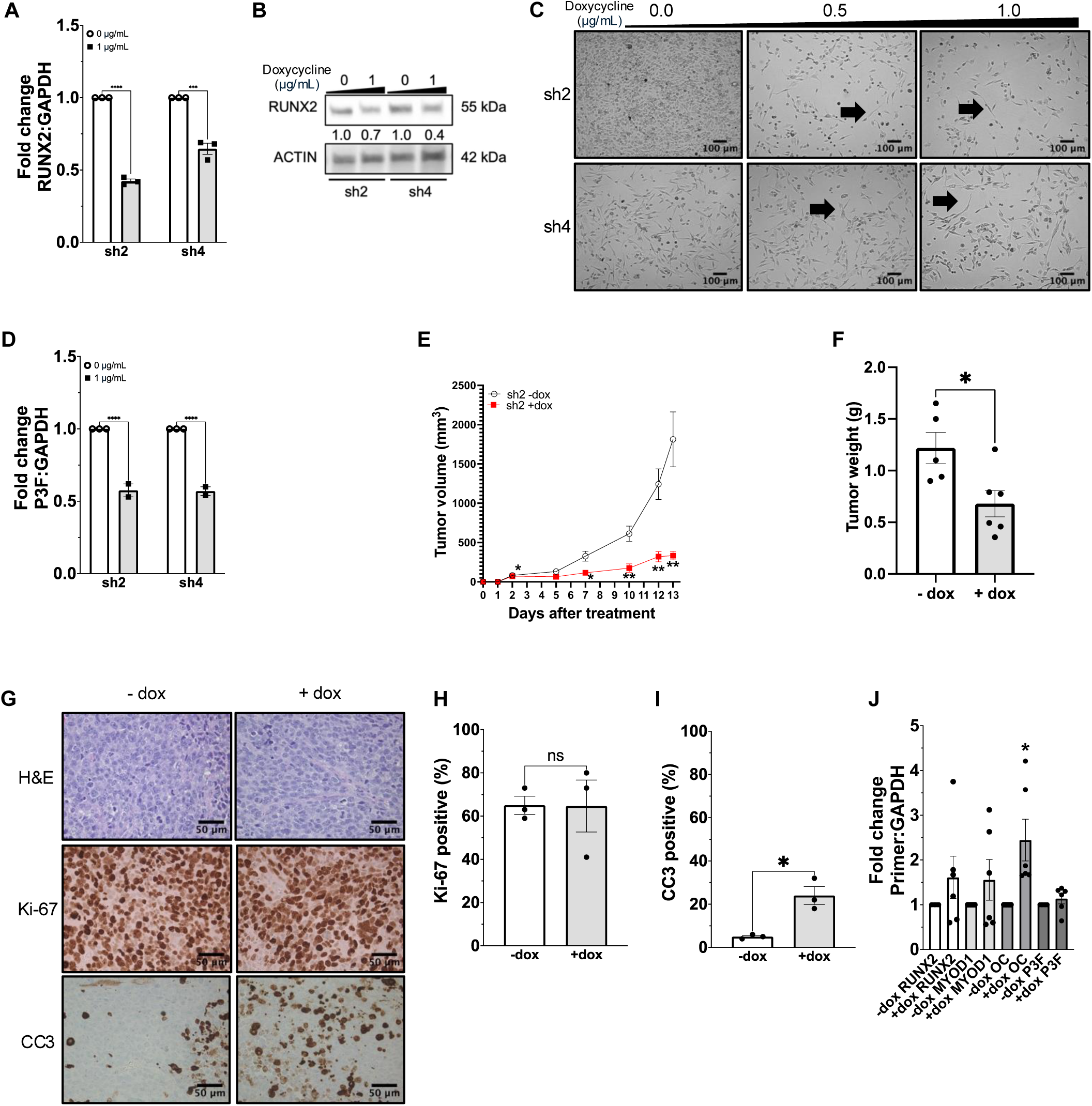
Doxycycline inducible *RUNX2* knockdown in Rh30 FP-RMS cells. **A,** qRT-PCR for *RUNX2* 72 h after 1 µg/mL doxycycline treatment. Black dots represent technical replicates. **B,** RUNX2 protein expression in *RUNX2* knockdown cells (sh2, sh4) following treatment with 1 µg/mL doxycycline for 140 h. **C,** Doxycycline inducible knockdown of *RUNX2* in Rh30 cells using two different shRNA constructs (sh2, sh4). Black arrows point to visible differentiated phenotypes. Cells were imaged at 10X magnification. Scale bars are 100 µm. **D,** qRT-PCR for *PAX3::FOXO1* expression 72 h after 1 µg/mL doxycycline treatment in the *RUNX2* knockdown cells (sh2, sh4). Black dots are technical replicates. **E,** Tumor volume growth curves in mice engrafted with sh2 doxycycline inducible *RUNX2* knockdown Rh30 cells and with or without access to doxycycline chow. Stars above each day represent a significant difference between means of control versus treated tumors on that day. **F,** Tumors excised from the mice and weighed (g). Black dots represent independent biological replicates. **G,** Representative H&E images of excised tumors taken at 40X magnification plus immunohistochemistry for Ki-67 and CC3. Scale bars are 50 µm. **H,I,** Three representative tumors from each group (one shown) stained for Ki-67 and CC3 and quantified using ImageJ. **J,** qRT-PCR for *RUNX2, MYOD1, OC,* and *PAX3::FOXO1* expression following *RUNX2* knockdown *in vivo*. Blak dots represent independent biological replicates.

### Identification of a feed-forward loop between PAX3::FOXO1 and RUNX2

Since *RUNX2* KD altered expression of *PAX3::FOXO1* and its target genes, we next evaluated whether *PAX3::FOXO1* conversely regulated *RUNX2.* Mining of previously published mRNA-seq datasets showed that *RUNX2* but not *RUNX1* expression is decreased following lentiviral KD of *PAX3::FOXO1* in Rh4 FP-RMS cells (**Fig. 5A,B**) (8). Prior studies in Rh30 cells showed that *RUNX2* was rapidly inhibited following PAX3::FOXO1 degradation (10). To expand *PAX3::FOXO1* KD findings, we transiently suppressed *PAX3::FOXO1* using siRNAs, and found that *PAX3::FOXO1* KD inhibited *RUNX2* expression by approximately 85% in Rh30 cells (**Fig. 5C,D**). Taken together, these data suggest that RUNX2 and PAX3::FOXO1 can regulate one another (**Fig. 4D**; **Fig. 5C,D**). This relationship is supported by the direct binding of PAX3::FOXO1 to *RUNX2* enhancer as determined by a previously published ChIP-seq dataset in Rh4 cells (**Fig. 5E**) (8). Previous studies reported similar results in Rh30 cells using CUT&RUN (10), thus collectively these data suggest that PAX3::FOXO1 binding to the *RUNX2* enhancer is not cell line-specific but likely occurs in all PAX3::FOXO1 expressing cell lines (8,10). Additional ChIP-seq analyses reveal active histone marks (H3K27ac, H3K9ac, and H3K4me3) but inactive repressive marks (H3K27me3) at the RUNX2 promoter, confirming that the *RUNX2* promoter is active when PAX3::FOXO1 is bound to the *RUNX2* enhancer (**Fig. 5E**). The core regulatory circuit of PAX3::FOXO1 associated transcription factors regulate each other and finding MYOD1 at the enhancer of *RUNX2* may indicate that *RUNX2* is an important target of the core regulatory circuit (**Fig. 5E**); since *MYOD1* increases when *RUNX2* is KD, it is unlikely that RUNX2 binds *MYOD1* in FP-RMS cells (**Fig. 2G; Supplementary TableS7**). This finding was validated in an additional cell line and expanded on in the previously published Rh30 CUT&RUN dataset where other PAX3::FOXO1 complex proteins such as PAX3, CDK8, RUNX1, and BRD4 localize at the same enhancer alongside PAX3::FOXO1 (10). Additional tracks visualizing ATAC-seq and mRNA-seq data demonstrates that the *RUNX2* gene is open and expressed in the presence of PAX3::FOXO1, but lost with *PAX3::FOXO1* KD; potentially because of the ability of PAX3::FOXO1 to coordinate enhancer architecture and remodel chromatin of enhancers and super enhancers (8,10).

**Figure 5.**
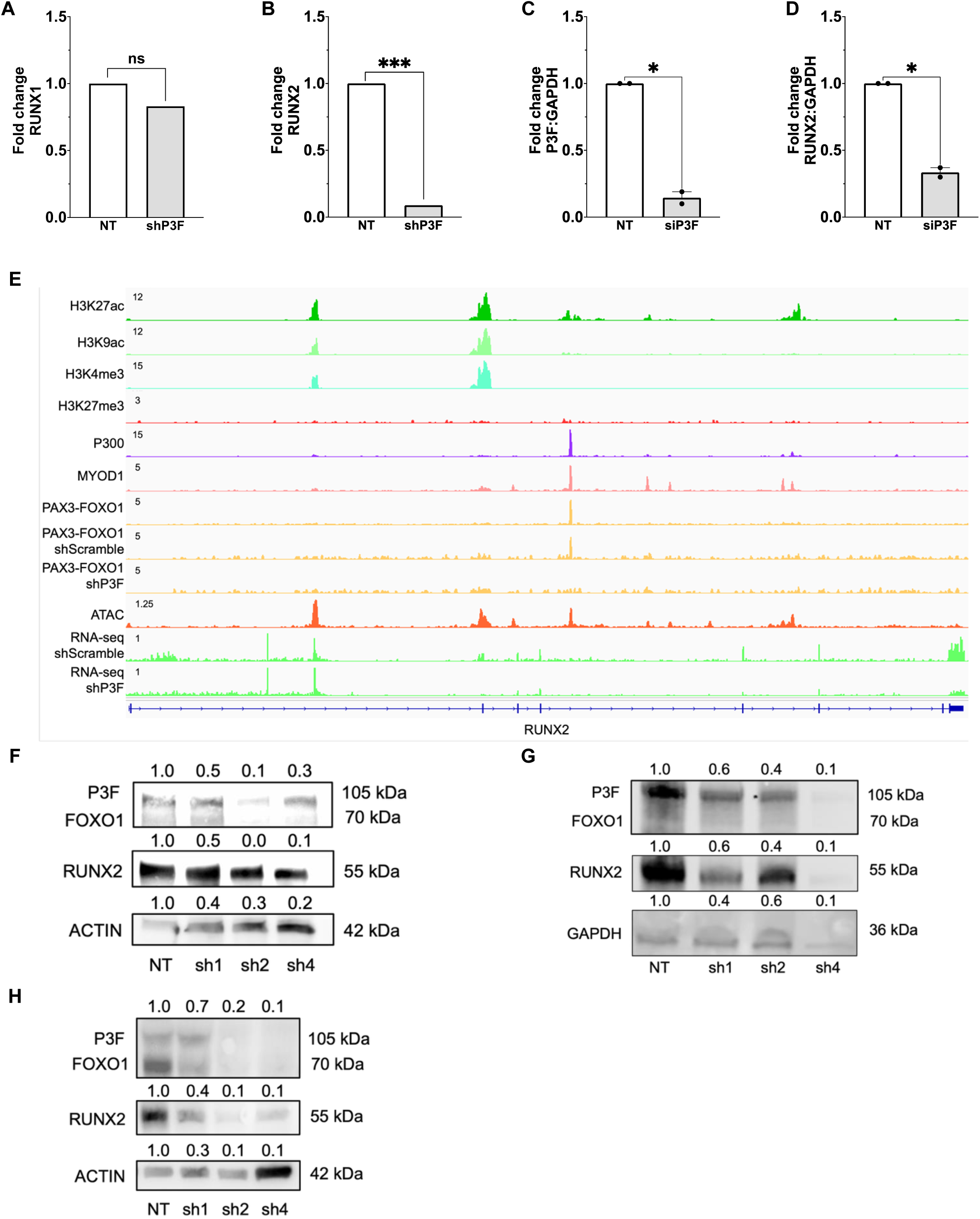
PAX3::FOXO1 and RUNX2 reciprocally regulate one another in FP-RMS. **A,B,** Stable knockdown of *PAX3*::*FOXO1* in Rh4 cells affect on *RUNX2* and *RUNX1* expression. **C,D,** qRT-PCR for *PAX3::FOXO1* and *RUNX2* expression in *PAX3::FOXO1* transient knockdown Rh30 cells. Black dots are independent biological replicates. **E,** Genome browser view of *RUNX2*. Peaks for the top 9 rows reflect degree of DNA-binding of H3K27ac, H3K9ac, H3K4me3, H3K27me3, P300, MYOD1, and PAX3::FOXO1 using ChIP-seq. Row 10 is open chromatin peaks from ATAC-seq, and rows 11 and 12 are mRNA-seq reads. All rows are in Rh4 cells, counts are in reads per million. **F,** PAX3::FOXO1, FOXO1, and RUNX2 protein expression in negative control (NT) and puromycin selected *RUNX2* knockdown Rh4 cells after 48 h selection. **G,** PAX3::FOXO1, FOXO1 and RUNX2 protein expression in negative control (NT) and puromycin selected *RUNX2* knockdown Rh30 cells after 72 h selection. **H,** PAX3::FOXO1, FOXO1 and RUNX2 protein expression in negative control (NT) and puromycin selected *RUNX2* knockdown Rh30 cells 144 h after selection.

Given the regulatory effects of PAX3::FOXO1 on *RUNX2,* and RUNX2 on *PAX3::FOXO1* and associated target genes, we investigated whether *RUNX2* KD impacted PAX3::FOXO1 protein expression. Indeed, inhibition of *RUNX2* resulted in decreased expression of PAX3::FOXO1 and endogenous FOXO1 in both Rh4 and Rh30 cells, signifying a novel mechanism wherein PAX3::FOXO1 protein and gene expression is dependent on the expression of RUNX2 (**Fig. 5F,G**). With longer RUNX2 KD (144 h after selection), PAX3::FOXO1 and endogenous FOXO1 expression were almost completely absent (**Fig. 5H**). Together, these findings establish a feed-forward loop between RUNX2 and PAX3::FOXO1 that regulates FP-RMS oncogenic phenotypes and regulatory gene networks.

### Pharmacologic inhibition of RUNX2 via the small molecule RUNX2 inhibitor CADD522 impairs FP-RMS tumor xenograft formation

To evaluate RUNX2 inhibition for clinical purposes, we investigated the efficacy of computer aided drug design molecule (CADD522), a commercially available small-molecule RUNX2 inhibitor previously tested pre-clinically in human breast cancer, Ewing sarcoma, and OS animal models (31,38,39,48), but not in FP-RMS models. To capture early genetic changes in response to CADD522 treatment, qRT-PCR of *RUNX2* significantly decreased with 1.0 μM CADD522 treatment for 24 h (**Fig. 6A**). Further *in vitro* investigation determined that treatment with 12 μM CADD522, similar to *RUNX2* KD, resulted in decreased colony number (**Fig. 6B,C**). Rh30 cells with 1.5 μM and 3.0 μM CADD522 for 72 h exhibited that drug treatment morphologically phenocopied genetic *RUNX2* KD (**Fig. 6D, top row**). Next, we performed MF20 staining on Rh30 cells and found that CADD522 treatment induced myotubes, thus supporting RUNX2’s effects on FP-RMS cell differentiation (**Fig. 6D, bottom row, E**). qRT-PCR of *MYOD1,* and *PAX3::FOXO1* partially mimicked results seen with *RUNX2* KD in a dose-dependent manner, but *OC* expression did not change with 0.5 μM and 1.0 μM CADD522 treatment for 24 h suggesting mechanistic differences between the drug and KD, likely because of differing mechanisms of inhibition and off target effects on metabolism (**Fig. 6F-H**) (38).

**Figure 6.**
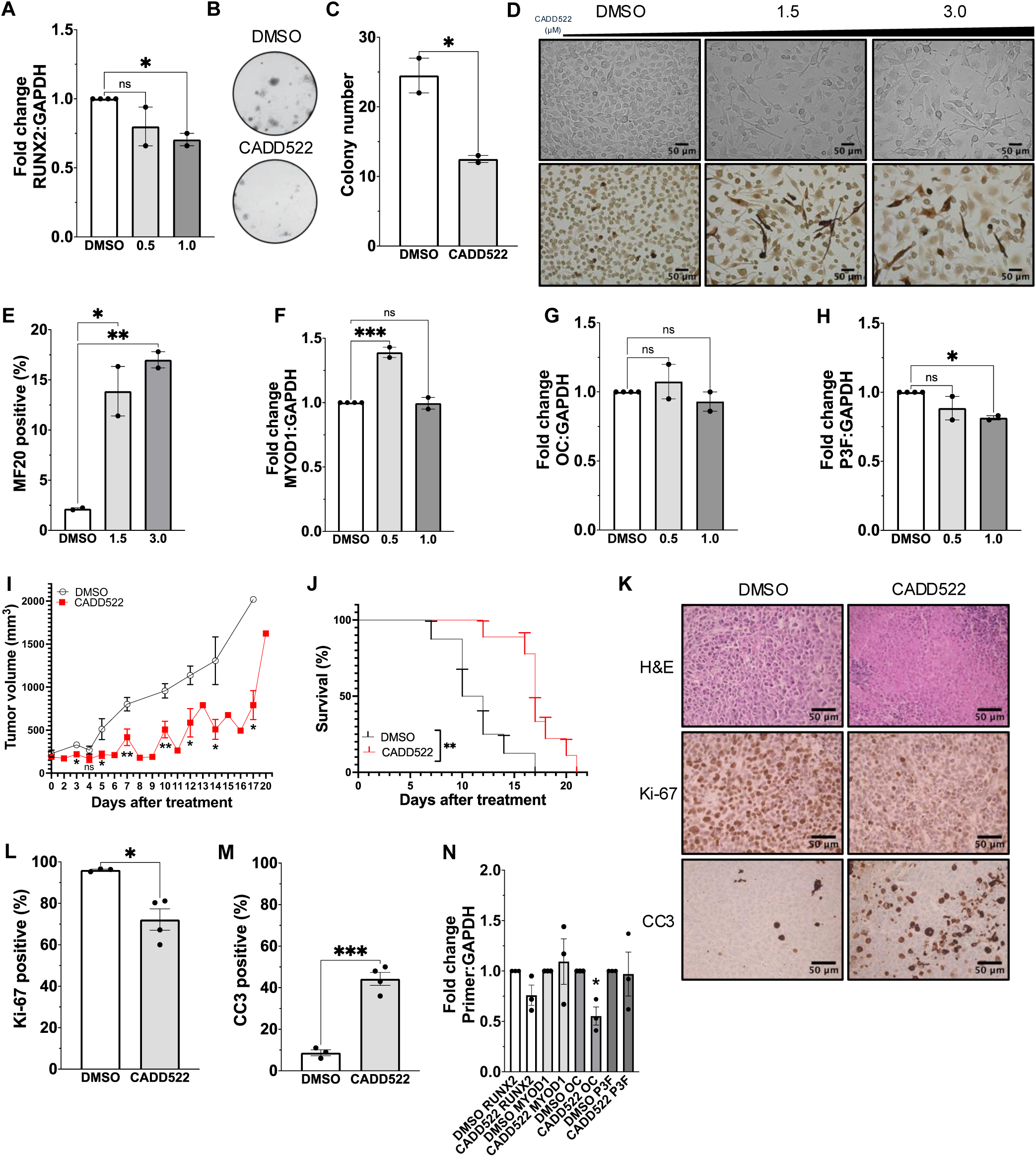
Rh30 cells treated with the RUNX2 small molecule inhibitor CADD522 *in vitro* and *in vivo.* **A,** qRT-PCR for *RUNX2* performed following 24 h of treatment with 0.5 μM and 1.0 μM of CADD522 *in vitro*. Black dots are independent biological replicates. **B,** CADD522 treatment effects on colony formation *in vitro*. Cells were treated with a 12 μM dose every other day for 14 d. **C,** Colony number after CADD522 treatment. Black dots represent technical replicates. **D,** Rh30 cells stained with the myogenic marker sarcomere myosin (MF20) when treated with a negative control (DMSO) and different CADD522 concentrations over 72 h. Images were captured at 20X magnification. Scale bars are 50 µm. **E,** Quantification of MF20 positive cells (dark brown) versus MF20 negative cells (light brown) following CADD522 treatment. **F, G, H** qRT-PCR for *MYOD1, OC,* and *PAX3::FOXO1* performed following 24 h of treatment with 0.5 μM and 1.0 μM of CADD522 *in vitro*. Black dots are independent biological replicates. **I,** Tumor volume growth curves with (n=9) or without 10 mg/kg CADD522 (n=8). **J,** Kaplan-Meier survival curve for mice treated with 10 mg/kg CADD522. Stars above each day represent a significant difference between means of control versus treated tumors on that day. **K,** H&E stains on excised tumors and immunohistochemistry (IHC) for Ki-67 and CC3 with and without CADD522 treatment. Images were captured at 40X magnification. Scale bars are 50 µm. **L,M,** Ki-67 and CC3 quantification. **N,** qRT-PCR for *RUNX2, MYOD1, OC,* and *PAX3::FOXO1* expression following CADD522 treatment *in vivo*. Black dots represent independent biological replicates

Expanded *in vitro* studies using different doses of and exposure to CADD522 are warranted to fully elucidate its mechanism in FP-RMS. Finally, to determine whether CADD522 is effective at suppressing tumor growth and extending survival *in vivo*, we injected Rh30 cells subcutaneously into the flanks of SCID/*beige* mice and treated them with 10 mg/kg of drug 3x weekly once tumors were 200 mm^3^. Mice in the drug treatment group had significantly increased survival and reduced tumor volume (**Fig. 6I; Supp. Fig. S5D**). Mice treated with CADD522 also had significantly increased survival (*p*=0.0017) (**Fig. 6J**). To characterize tumors further, we performed IHC for Ki-67 and CC3 and determined that drug treated tumors were significantly less proliferative and significantly more apoptotic (**Fig. 6K,I**). Finally, qRT-PCR for *RUNX2, MYOD1 OC,* and *PAX3::FOXO1* closely resembled *in vitro* findings seen in **Fig. 2** and **Fig. 4**, except for *PAX3::FOXO1* expression, which was variable, and *OC* expression, which was significantly decreased, suggesting that dose optimization and continue studies are required to fully determine the mechanistic effects of CADD522 in FP-RMS (**Fig. 6N**).

### microRNA mediated regulation of RUNX2 in ARMS patient samples

Since RUNX2 expression is also known to be regulated by microRNAs (miRNAs), specifically miR-34a (49), in parallel with our mRNA-seq of primary tumor tissue, we also performed sRNA-seq, to identify novel and validate previously published miRNAs responsible for regulating key RMS genes. Representative frequency mapping of miRNA expression in wild-type and RMS tissues revealed that SkM tissues predominantly expressed miRNAs at 22 nt, while RMS tissues exhibited two major peaks: one at 22 nt for miRNAs and another at 32 nt for tRNA-derived fragments (tRFs) and Y-RNA small RNA (ysRNAs) (**Supp. Fig. S5A**). The PCC plot demonstrated a strong linear correlation between the sample sets, indicating consistency within each group (**Supp. Fig. S5B**). PCA further supported these findings, showing distinct clustering along the PC1 axis that corresponded to ARMS, ERMS, and SkM tissues (**Supp. Fig. S5C**). Approximately 15M reads per sample were obtained (**Supplementary Table S1**). Hierarchical cluster analysis of selected miRNAs in ARMS tissues compared to SkM revealed significant changes in miRNA expression, providing insight into their role in regulating RMS biology (**Supp. Fig. S5D**). Similarly, a heatmap of selected miRNAs across ERMS tissues compared to SkM highlighted additional miRNAs involved in RMS regulation (**Fig. S5E**). Processed data for all miRNAs can be found in **Supplementary Table S9.** miR-34a was identified as highly expressed in all RMS histological subtypes coinciding with its known role of supporting *RUNX2* expression in patients with calcific aortic valve stenosis (**Supplementary Table S9**) (31). These data suggest that RUNX2 is likely regulated by specific miRNAs in FP-RMS, some of which may make interesting future therapeutic targets, and be critical to RUNX2-PAX3::FOXO1 signaling in FP-RMS.

## Discussion

RMS is the most common soft tissue sarcoma diagnosed in the pediatric population (1–3,5). Prior studies have revealed that around 1 in 4 RMS cases harbor a *PAX3/7*::*FOXO1* driver mutation (1–3,5). The encoded fusion oncoproteins create *de novo* enhancers (8,10) that appear to contribute to tumorigenesis and eventually tumor progression; considered high-risk disease with higher chemoresistance and increased metastatic propensity (2,5,50). The PAX3/7::FOXO1 oncoproteins are ideal targets for cancer drugs because they are unique to RMS cells; however, they possess no catalytic activity and are intrinsically disordered proteins with ill-defined binding pockets for small molecule inhibitors, meaning alternative strategies are required to target RMS cells. Here we perform sequencing of pediatric primary tumor samples and cross reference publicly available datasets to identify RUNX2, long considered a bone development transcription factor, as being critical for the oncogenic properties of FP-RMS, a cancer more associated with the skeletal muscle lineage. Short read sequencing of patient tumors was employed for differential expressed gene analysis using the original histological diagnosis, future work will employ long read sequencing and diagnosis based on fusion status going forward, in combination with short read (Illumina) sequencing. Data mining from non-malignant tissue and developmental pathways indeed places RUNX2 squarely in myogenic processes. For example, GTExPortal snRNA-seq data of skeletal muscle tissue revealed that RUNX2 is expressed in 2.43% of skeletal muscle myocyte cells (**Supp. Fig. S2A**), supporting its context-specific function in normal tissue. There is evidence that RUNX2 is critical for cell fate decisions (e.g. bone and muscle lineages), since some muscle progenitor cells are bipotent and can be directed towards osteogenic or myogenic lineages depending upon modifications by lysine methyltransferases (LSD1) or lysine demethylases (MLL4), which have opposing functions at *RUNX2* enhancers to regulate myogenic versus osteogenic differentiation (18). While the relationship between PAX3::FOXO1 and RUNX2 is largely unknown, several studies have interrogated the relationship between FOXO1 and RUNX2 in other contexts (49,51).

In prostate cancer, *RUNX2* governed migration and invasion is negatively regulated by increased *FOXO1* expression; in early osteoblast differentiation, amino acids 360-456 of FOXO1 directly interacts with amino acids 242-258 of RUNX2 to dissociate RUNX2 from the *osteocalcin* (*OC*) promoter (49,51). In the case of FP-RMS, it is likely that RUNX2 upregulation contributes to the core regulatory circuitry maintaining the cells in a myoblastic state, since when we suppress *RUNX2,* genes responsible for myogenesis are upregulated, while genes for cell cycle inhibition and PAX3::FOXO1 specific gene signatures meditating the anti-apoptotic function of PAX3::FOXO1 (50) are downregulated (**Fig. 7**). Mechanistically, this appears due to a feed-forward loop, whereby PAX3::FOXO1 transcriptionally upregulates RUNX2, and RUNX2 maintains PAX3::FOXO1 transcript levels, sustaining PAX3::FOXO1 signaling to genes such as *MYCN, POU4F1, RASSF4,* and *FGFR2.* At the chromatin level, it is more complex, since when a PAX3::FOXO1 regulated-enhancer of RUNX2 was deleted following depletion of PAX3::FOXO1, *RUNX2* expression decreased, but when the non-PAX3::FOXO1 enhancer was deleted, the effect was the same; therefore, two elements are independently required to maintain *RUNX2* gene expression (10). These data suggest that multiple mechanisms likely regulate RUNX2 in FP-RMS, some of which may be pertinent to FN-RMS. For example, single cell transcriptomic profiling of FN-RMS cells identified a molecularly-defined tumor-propagating subpopulation that shares similarity to bipotent, MSCs capable of differentiating into muscle and osteogenic cells (44). These observations, in combination with our finding that *RUNX2* expression is high not only in FP-RMS but also in histologically defined ERMS, SCRMS, and PRMS patient samples (strongly associated with FN-RMS) suggest that RUNX2 oncogenic signaling may be relevant beyond its relationship with PAX3::FOXO1 and FP-RMS (50), and may be a pan-RMS target.

**Figure 7.**
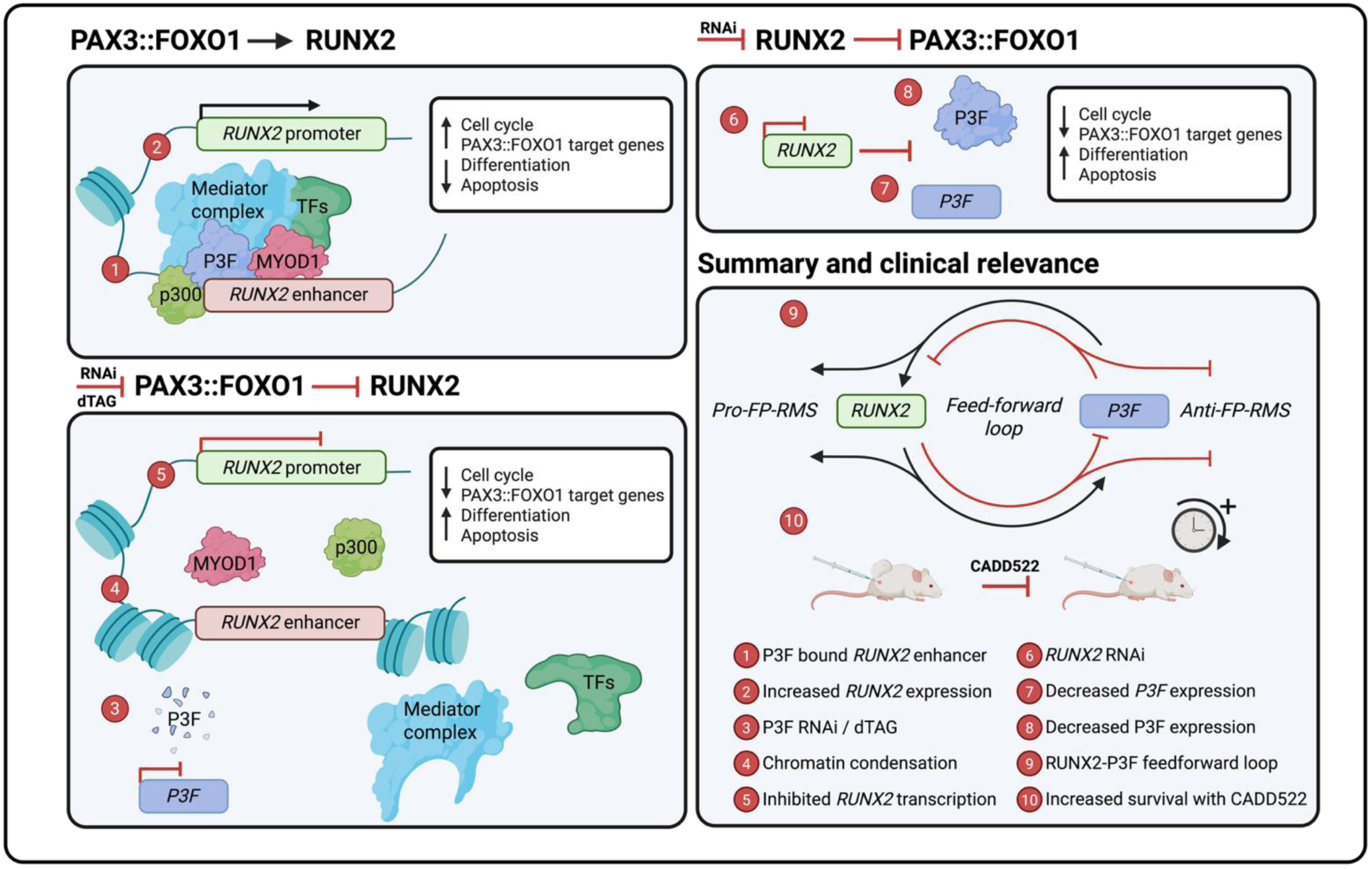
Schematic to show PAX3::FOXO1 transcriptionally upregulates *RUNX2* expression at the chromatin to prevent FP-RMS terminal differentiation and apoptosis via the defined PAX3::FOXO1 core regulatory circuitry.

A relatively unexplored but intriguing direction supported by our work is the relationship of RUNX1 to RUNX2 in FP-RMS. In non-cancer situations, cooperation of RUNX1 and RUNX2 are required to regulate sternal morphogenesis and the commitment of MSCs to chondrocytes (15,16,52). Several platforms and the literature highlight RUNX1 as a prognostic biomarker across many human malignancies including hepatocellular carcinoma and gastric cancer. On the other hand, RUNX1 is tumor-suppressive in lung and prostate cancer (53). In breast cancer stem cells, RUNX1 and RUNX2 serve opposing roles (52). Germline mutations of *RUNX1* are associated with myeloid malignancies, and *RUNX1* is required for growth and survival of multiple leukemia cell types (54). Therefore, it is likely that the behaviors and contributions of RUNX1 and RUNX2 to cancer is context-dependent. Here, we found that *RUNX1* is upregulated in ERMS but is not differentially expressed in ARMS (ERMS: log_2_FC=1.4); survival data suggests that low *RUNX1* expression is correlated with poor outcome (8). Regarding mechanism, there are conflicting reports on whether *RUNX1* is transcriptionally regulated by PAX3::FOXO1. Through analysis of published datasets, we found that shRNA KD of *PAX3::FOXO1* did not affect *RUNX1* expression, and that PAX3::FOXO1 did not bind *RUNX1* enhancers (8). This finding may in part explain why PAX3::FOXO1 was found to regulate RUNX1 enhancer independently, but regulated RUNX2 enhancer alongside other DNA-binding factors such as MYOD1 (10). However, RUNX1 is part of the PAX3::FOXO1 protein interactome; and that *RUNX1* is downregulated upon loss of PAX3::FOXO1 protein within 6 h of dTAG-47 treatment (10). These data may be of significance for the future understanding of the role of RUNX1 in FP-RMS and deciphering a potentially coregulatory relationship between RUNX1 and RUNX2 in FP-RMS.

The clinical relevance of RUNX2 rests on its use as a biomarker of outcome and as a therapeutic target. Because our sample size was small, it is too early to say whether RUNX2 is a bona fide biomarker in FP-RMS. RUNX2 is a prognostic cancer biomarker in diseases such as OS, blastic plasmacytoid dendritic cell neoplasm (BPDCN), breast cancer, and prostate cancer; indirect and direct biological functions of RUNX2 regulate oncogenic processes such as cancer stemness, metastasis, proliferation, cell cycle regulation, and apoptosis (31,39,55–59). In BPDCN, the plasmacytoid dendritic cells (pDCs) *RUNX2* super-enhancer is hijacked to activate *MYC* in and *RUNX2* expression to promote proliferation (57).

However, given the slowing of tumor xenograft growth by either genetic or CADD522 pharmacologic inhibition of RUNX2, the value of RUNX2 as a therapeutic target should be pursued. Transcription factors including RUNX2 remain challenging as drug targets because of their lack of catalytic activity and their intrinsically disordered state, lacking binding pockets for small molecules. However, the RUNX2-directed agent used here, CADD522, is in development for clinical trials (60), and others are developing allosteric inhibitors to RUNX2 and CBFβ inhibitors (61). Like CADD522, active inhibitor compounds demonstrate anti-tumor activities with concentrations that are achievable *in vivo* (31,39,61). There may also be indirect ways to target RUNX1/RUNX2. The small molecule imatinib, FDA-approved for the treatment of multiple hematologic malignancies, is being examined in clinical trials as a means to block tyrosine phosphorylation of RUNX1, thereby restoring RUNX1 activity, especially in patients with RUNX1 germline mutations (62,63). These data highlight the importance of RUNX-targeted therapies across various cancer types and the demand for other RUNX-targeted pharmacologic approaches such as PROTACS and molecular glues.

Finally, we query what other pathways may control RUNX2 expression, since targets in these pathways may also be required to keep RUNX2 expression high. For example, HDAC4, which is a direct negative regulator of *RUNX2* promoter accessibility (64), was downregulated in both ARMS (log_2_FC=1.1) and ERMS (log_2_FC=2.14) samples, which might underlie the observed *RUNX2* upregulation. Because microRNAs are known to regulate HDAC4, we performed sRNA-seq using the same samples subject to mRNA profiling. Most of the miRNAs known to negatively regulate *HDAC4* – therefore permitting expression of HDAC4 targets including *RUNX2* - were paradoxically downregulated [e.g. miR-140 (both 3p and 5p isoforms), muscle specific miR-1 and miR-206] except for miR-155, which was upregulated (ARMS: log_2_FC=2.0; ERMS: log_2_FC=3.1). A future study could directly test a miR-155 - HDAC4 - RUNX2 axis in RMS because miR-155 was previously reported as a poor prognostic marker in sarcoma; the current data might explain the causal mechanism for the previously reported observation (65). To provide future directions, all differentially expressed microRNAs were funneled through the DepMap (**Supp. Fig. S6F**) to identify microRNAs exclusively shared between all RMS histological subtypes (43). The 81 exclusively shared microRNAs, which included miR-140 isoforms and miR-155, can be found in **Supplementary Table S9.**

In summary, here we have corroborated the upregulation of RUNX2 in FP-RMS and other RMS variants and newly identified a feed-forward loop, whereby PAX3::FOXO1 transcriptionally upregulates RUNX2, and RUNX2 supports the expression of PAX3::FOXO1. We have demonstrated that loss of RUNX2 in FP-RMS abrogates oncogenic phenotypes *in vitro*, and that genetic or pharmacologic blockade of RUNX2 *in vivo* impairs FP-RMS tumor growth. Future studies will focus on the development of RUNX2 as therapeutic target and the molecular mechanisms of RUNX2 regulation.

## Supporting information

Table S5C RMS dependencies Zhang et al

Table S5D RMS dependencies Gryder et al

Table S6 QC mRNS RUNX2 KD

Table S7. DE_RH4_sh4_0h_vs_RH4_shNT_0h

Table S7. DE_RH4_sh4_24h_vs_RH4_shNT_24h

Table S7. DE_RH4_sh4_48h_vs_RH4_shNT_48h

Table S7. DE_RH30_sh4_0h_vs_RH30_shNT_0h

Table S7. DE_RH30_sh4_24h_vs_RH30_shNT_24h

Table S7. DE_RH30_sh4_48h_vs_RH30_shNT_48h

Table S8. GSEA mRNA RUNX2 KD C2

Table S8. GSEA mRNA RUNX2 KD C6

Table S8. GSEA mRNA RUNX2 KD C7

Table S8. GSEA mRNA RUNX2 KD Hallmarks 2

Table S8. GSEA mRNA RUNX2 KD Hallmarks

Table S8. GSEA mRNA RUNX2 KD NCI Geneset

Table S8. GSEA mRNA RUNX2 KD Top250

Table S9. Differential expression miRNA ARMS selected

Table S9. Differential expression miRNA ARMS

Table S9. Differential expression miRNA ERMS selected

Table S9. Differential expression miRNA ERMS

Table S1. Seq QC mRNA; miRNA

Table S2A. Primers and constructs

Table S2B. Primers and constructs

Table S2C. Primers and constructs

Table S3A. Differential expression mRNA ARMS

Table S3B. Differential expression mRNA ERMS

Table S4A. UpSet mRNA; miRNA ARMS

Table S4B. UpSet mRNA; miRNA ERMS

Table S5A. FPRMS v All Cancer dependencies

Table S5B. FPRMS dependencies

## Conflict of interest statement

CML’s spouse is founder and owner of Grid Therapeutics, which is developing a monoclonal antibody for adult lung cancer. CML’s lab has received funding from Ryvu. Neither of these are related to the research in this manuscript. DG reports patents EP3897609B1 and WO2023209077A1.

## Acknowledgements

We thank members of the NCI FusOnC2 group and the laboratories of Dr. Gerry Blobe, Dr. Tammara Watts, and Dr. Michael Deel (Duke) for helpful discussions. We thank Dr. Katherine Gonzales, Dr. Xiangfeng Shen, Dr. Edward O’Brien, and Dr. Manqi Zhang for assistance with IHC and imaging. We thank Dr. John Bushweller for participating in helpful conversations about RUNX2 pharmacological inhibition. We thank the Duke University BioRepository and Precision Pathology Center (Duke BRPC; supported by P30CA014236) and the NCI’s Cooperative Human Tissue Network (CHTN (RRID:SCR_004446); supported at Duke University by UM1CA239755 for processing histological tissues. CML’s lab and team was supported by NIH grant 1U54CA231630 (to CML), NIH grant 5T32GM142605-03 (supporting EAM), and a Duke OBGE Precision Genomics Collaboratory Grant (to EAM). SW was supported by the Pediatric Oncology Student Training (POST) Program. AM was supported by the CureSearch for Children’s Cancer Acceleration Initiative Award. AJ was supported by the CA230526 Department of Defense Grant. AC was supported by the NSF Computers in Science and Math Undergraduate Education award 1834159. DG’s lab and members, ME, AS were supported by Children with Cancer UK and donations made to the Childhood, Adolescent and Young Adult Cancer Research Programme at the University of East Anglia. The authors thank Dionne Wortley for support with obtaining patient samples. Fig. 1F, Fig. 2D, Fig. 7, and Supp. Fig.S6F were Created in BioRender (Mendes, E. (2025) https://BioRender.com/26c5ptj, https://BioRender.com/tk0t2im, https://BioRender.com/kggsksv).

---

**Figure S1.**
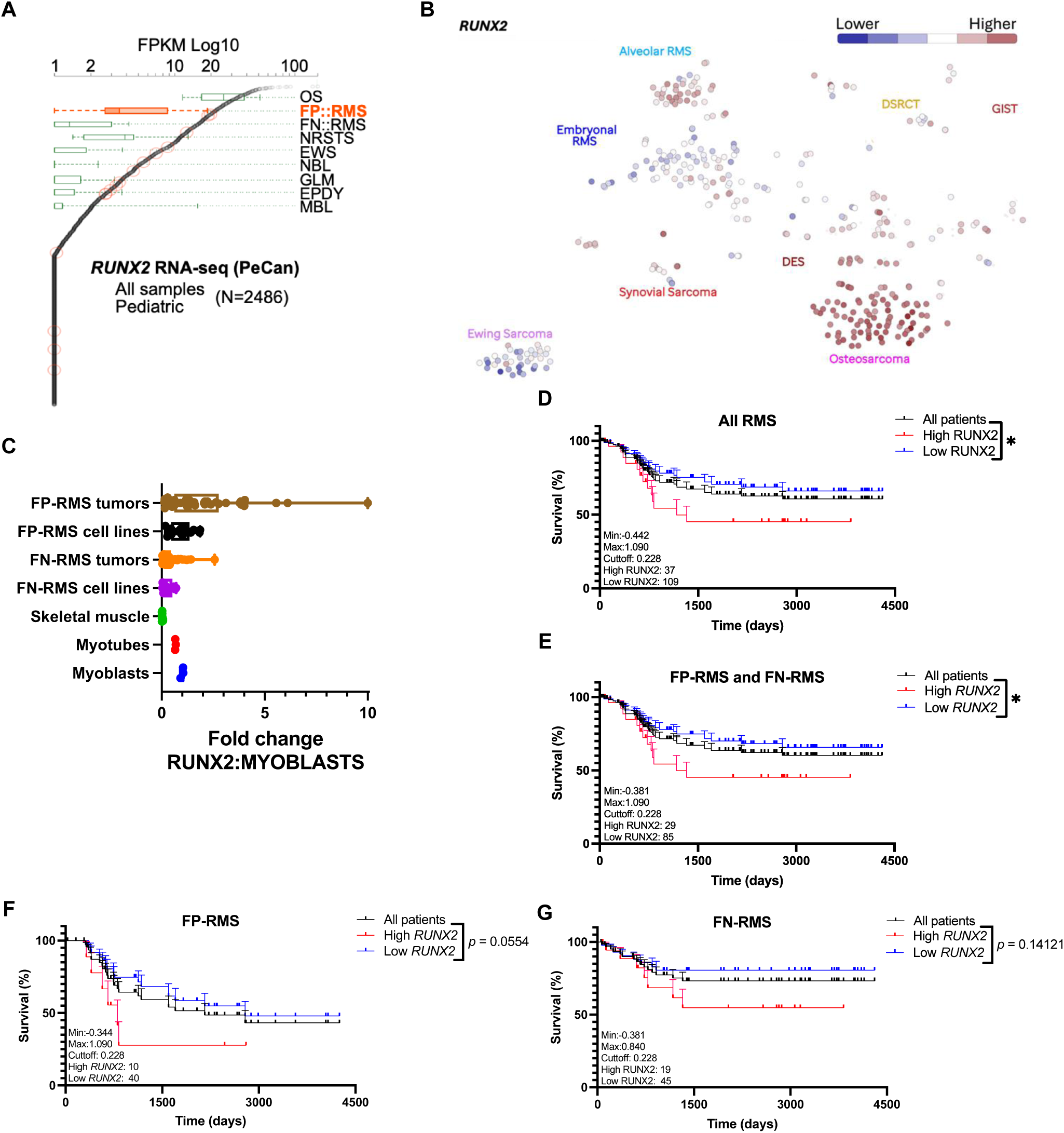
Expression and survival analysis of *RUNX2* in FP-RMS and FN-RMS. **A,** Analysis of *RUNX2* expression in a cohort of 2486 pediatric cancers showing expression of *RUNX2* in OS as well as FP-RMS and NRSTS compared to expression in other pediatric solid tumors. OS, osteosarcoma (n=112), FP-RMS, fusion-positive rhabdomyosarcoma (n=16), FN-RMS, fusion-negative rhabdomyosarcoma, NRSTS (n=28), non-rhabdomyosarcoma soft tissue sarcoma (n=13), EWS, Ewing sarcoma (n=5), NBL, neuroblastoma (n=198), GLM, glioblastoma (n=182), EPDY, ependymoma (n=92), MBL, medulloblastoma (n=31). Other pediatric solid tumors shown for comparison. **B,** Cluster plot of *RUNX2* expression in pediatric bone and soft tissue sarcomas. **C,** *RUNX2* expression in FP-RMS tumors compared to myoblasts. Myoblast (n=3), myotube (n=3), skeletal muscle (n=14), FN cell line (n=12), FN tumor (n=63), FP cell line (n=16), FP tumor (n=39). **D,E,F,G** Kaplan-Meier plot demonstrates the correlation between *RUNX2* and survival in all RMS (n=146), FP-RMS and FN-RMS combined (n=114), FP-RMS (n=50), and FN-RMS (n=64).

**Figure S2.**
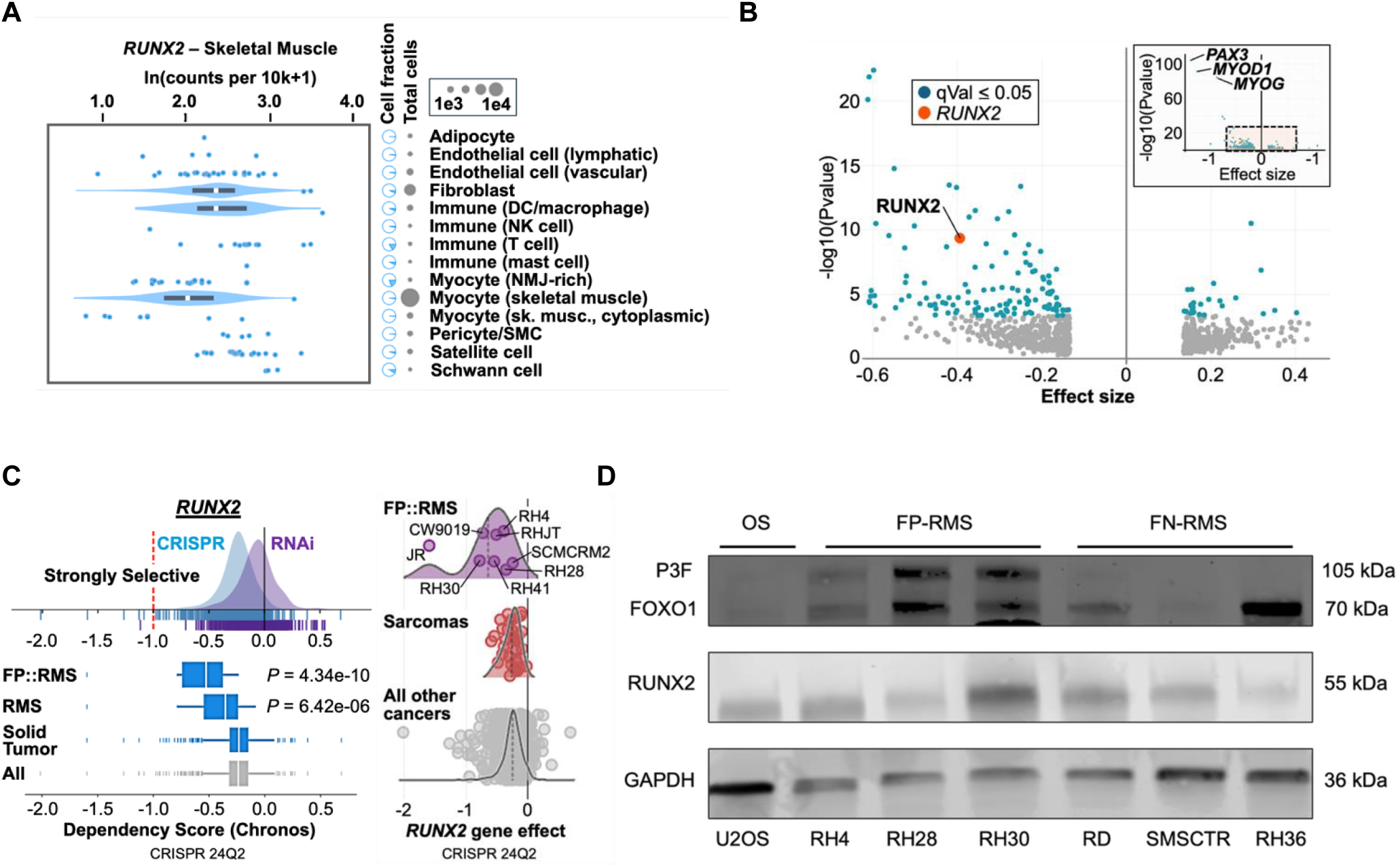
*RUNX2* dependency analysis and characterization in RMS. **A,** Expression of *RUNX2* in normal skeletal muscle. Skeletal muscle myocyte expressing *RUNX2* (n=504), total cells (n=20772). **B,** Volcano plot showing gene dependencies in FP-RMS cell lines relative to all other cancer cell lines. Box in upper right -hand corner are all the genes in the dataset and larger figure is a zoom in on the dashed box **Supplemental Table 5**. **C,** (Left side) Box plot of the CERES dependency score demonstrating *RUNX2* specific dependencies in FP-RMS compared to all cancer types analyzed together. FP-RMS (n=8), RMS (n=13), Solid Tumor (n=1019), All Cancer. (Right side) Characterization of *RUNX2* gene effect on FP-RMS cell lines compared to all sarcomas, and all other cancer. FP-RMS (n=8), RMS (n=13), Solid Tumor (n=1019), All Cancer. **D,** Immunoblot of PAX3::FOXO1, FOXO1, and RUNX2 protein expression in OS, FP-RMS, and FN-RMS cell lines.

**Figure S3.**
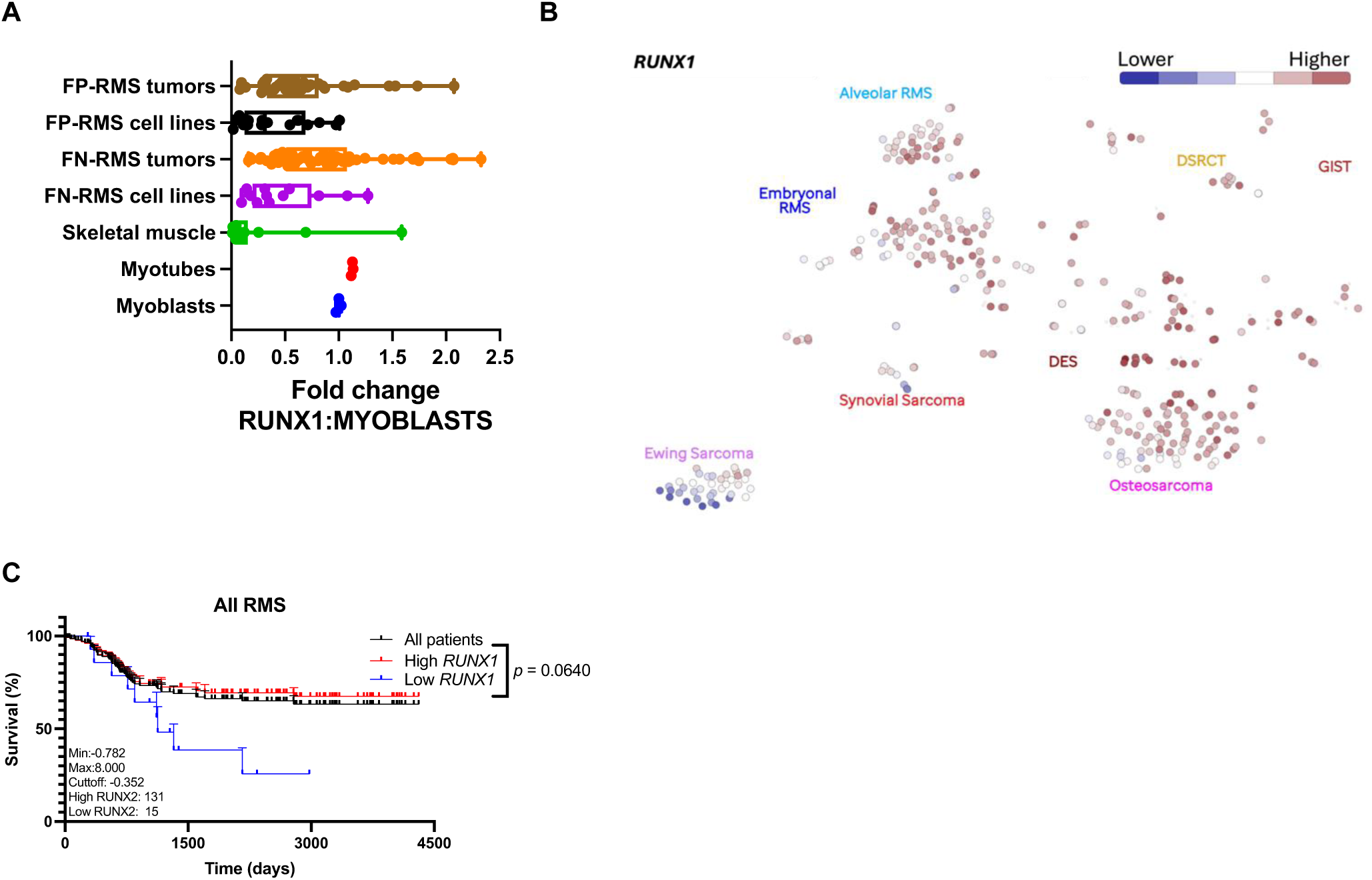
Expression and survival analysis of *RUNX1* in FP-RMS and FN-RMS. **A,** *RUNX1* expression in FN-RMS tumors and cell lines compared to myoblasts. Myoblast (n=3), myotube (n=3), skeletal muscle (n=14), FN cell line (n=12), FN tumor (n=63), FP cell line (n=16), FP tumor (n=39). **B,** Cluster plot of *RUNX1* expression in pediatric bone and soft tissue **C,** Kaplan-Meier plot demonstrates the correlation between *RUNX1* and survival in all RMS.

**Figure S4.**
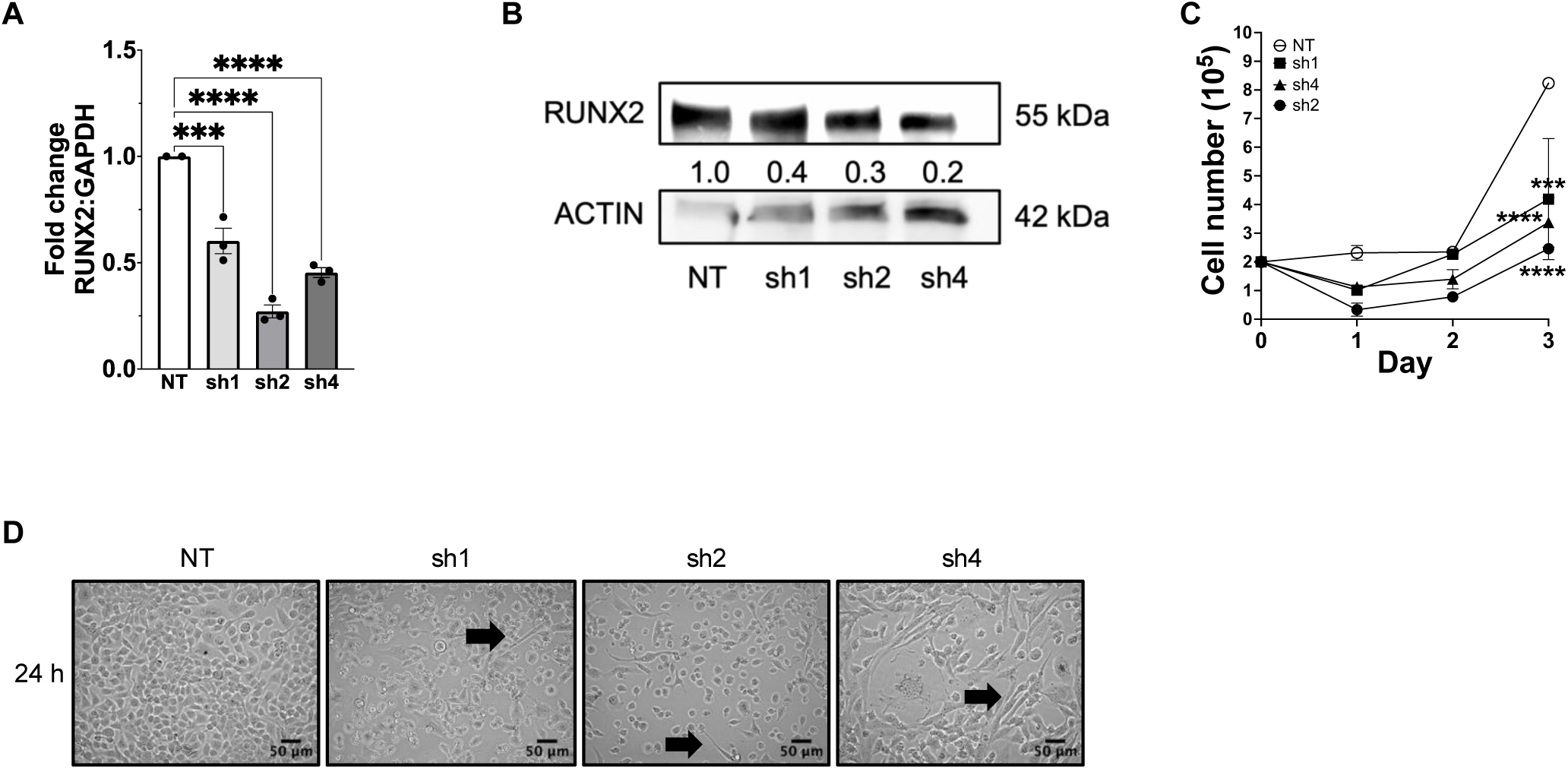
*RUNX2* knockdown in human FP-RMS cells (Rh4) impairs classical oncogenic phenotypes *in vitro.* **A,** qRT-PCR for *RUNX2* quantification in the negative control (NT) Rh4 cells and the three independent knockdown cells (sh1, sh2, sh4). Black dots are technical replicates. **B,** Normalized RUNX2 protein expression in the negative control (NT) Rh4 cells and the three independent knockdown cells (sh1, sh2, sh4) measured at 48 h. This blot is identical to Fig. 5F. **C,** Cell growth in negative control and *RUNX2* stable knockdown Rh30 cells. Black dots are independent biological replicates. **D,** *RUNX2* knockdown through three independent stably expressed shRUNX2 lentiviral preparations (sh1, sh2, sh4) plus negative control (shNT) in Rh4 cells. Black arrows point to visible areas of differentiation. Cells were imaged at 20X magnification and captured at 24 h. Scale bars are 50 µm.

**Figure S5.**
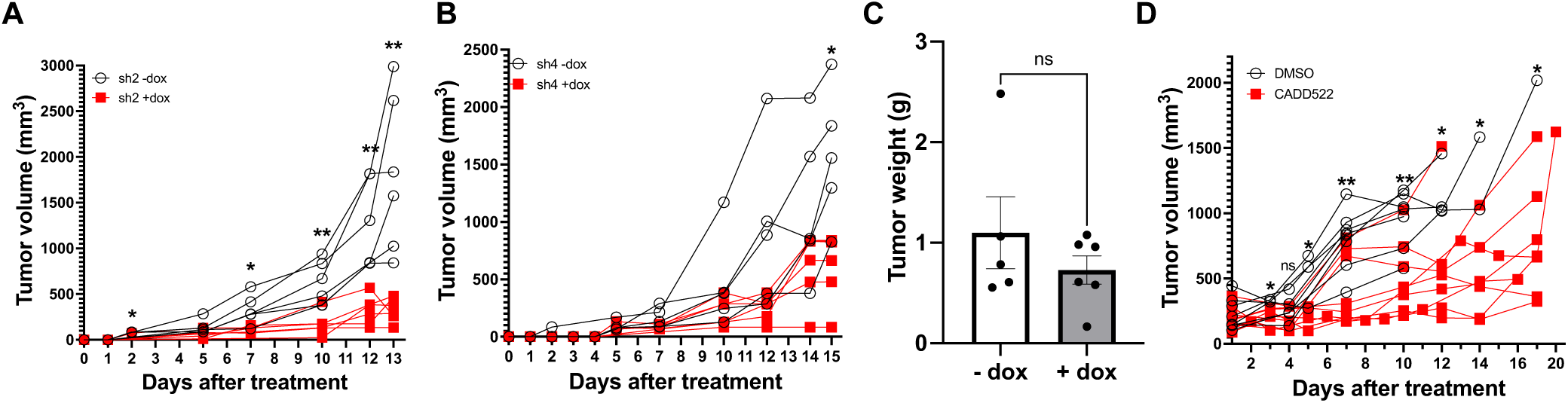
Additional *in vivo* experiments and expanded analyses. **A,** Tumor volume spider plot in mice engrafted with sh2 doxycycline inducible *RUNX2* knockdown Rh30 cells and with or without access to doxycycline chow (see Fig. 4). Each line represents a single mouse. **B,** Tumor volume spider plot in mice engrafted with sh4 doxycycline inducible *RUNX2* knockdown Rh30 cells and with or without access to doxycycline chow. Each line represents a single mouse. **C,** Tumors excised from the mice and weighed (g). Black dots represent independent biological replicates. **D,** Tumor volume spider plot in mice engrafted with wild type Rh30 cells with or without CADD522 treatment (see Fig. 6). Each line represents a single mouse. Stars above each day represent a significant difference between means of control versus treated tumors on that day.

**Figure S6.**
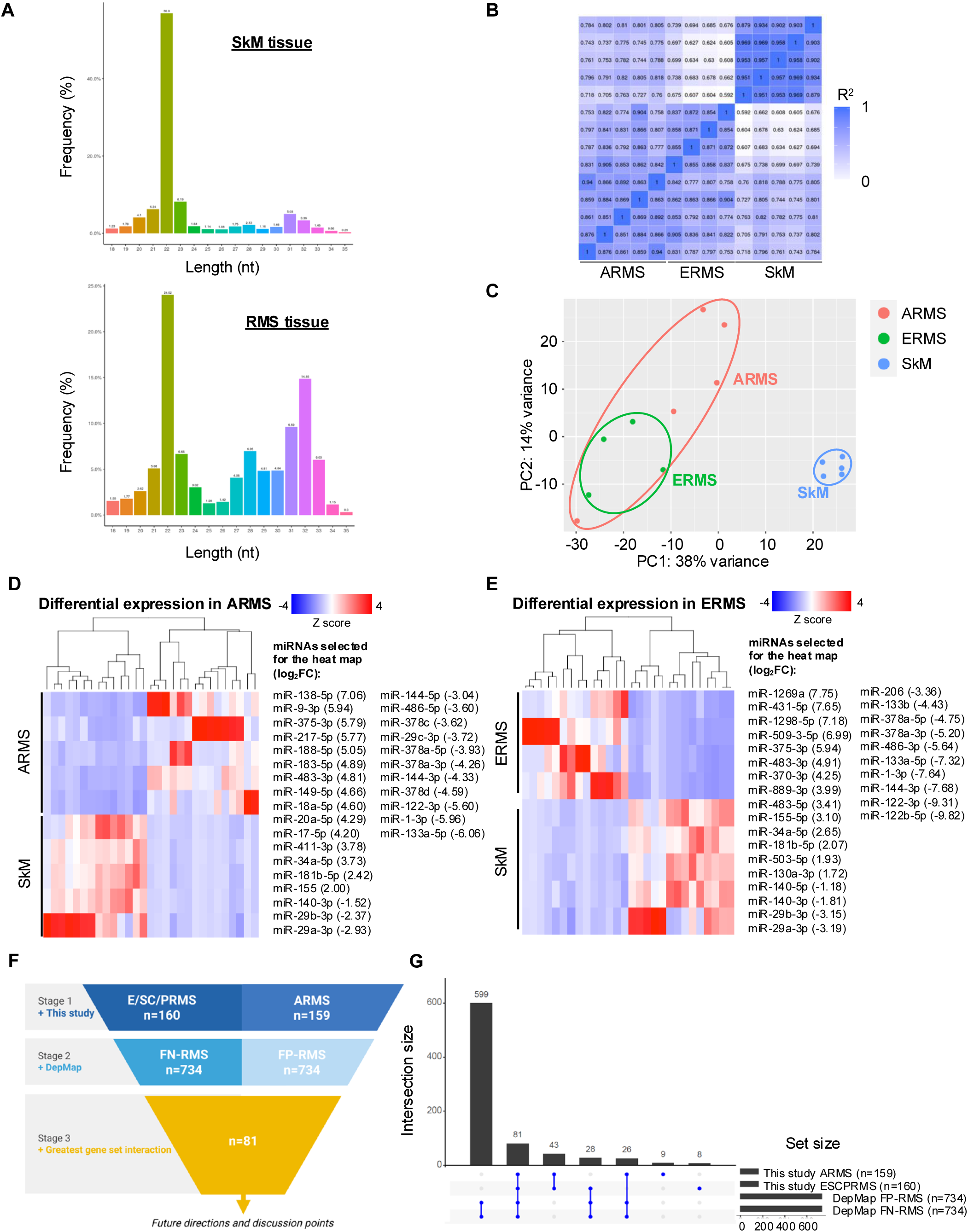
sRNA-seq performed on archival fresh frozen RMS tissues classified according to their histological interpretation. **A,** Representative frequency mapping graphs in SkM and RMS tissues shows that the SkM mostly expressed miRNAs at 22 nt. In RMS tissues, there were two major peaks: one at the miRNAs at 22 nt and one at tRFs and ysRNAs at 32 nt. **B,** Pearson correlation coefficient (PCC) plot shows a linear correlation between the sample sets. **C,** Data reduction to two-dimensions via biplot principal component analysis (PCA) shows consistent groups along the PC1 axis that correspond to ARMS (light red), ERMS (green) and SkM (blue) tissues. **D, E** Heat map based hierarchical cluster analysis of selected miRNAs (x-axis) across ARMS and ERMS tissues (y-axis) when compared to the SkM with the log_2_ fold change in expression also shown. Z-score refers to high (red) and low (blue) gene expression using normalized values when compared to the mean of total sequencing reads. **F,** Candidate selection of differentially expressed microRNAs across the current study (stage 1) and the cancer DepMap (stage 2). Candidate microRNAs of interest and discussion included those with the greatest gene set intersection (stage 3). Top candidate is bolded. **G,** UpSet plot to visualise intersected miRNAs (and therefore generate miRNA/s-of-interest for downstream studies) across the current study and the DepMap Portal.

## Notes

https://www.ncbi.nlm.nih.gov/geo/query/acc.cgi

### Competing Interest Statement

Spouse of CML is founder and owner of Grid Therapeutics, which is developing a monoclonal antibody for adult lung cancer. Lab of CML has received funding from Ryvu. Neither of these are related to the research in this manuscript. DG reports patents EP3897609B1 and WO2023209077A1.

## References

1. Linardic CM. PAX3-FOXO1 fusion gene in rhabdomyosarcoma. Cancer Lett 2008;270:10–8

2. Chen C, Dorado Garcia H, Scheer M, Henssen AG. Current and Future Treatment Strategies for Rhabdomyosarcoma. Front Oncol 2019;9:1458

3. Heske CM, Chi YY, Venkatramani R, Li M, Arnold MA, Dasgupta R, et al. Survival outcomes of patients with localized FOXO1 fusion-positive rhabdomyosarcoma treated on recent clinical trials: A report from the Soft Tissue Sarcoma Committee of the Children’s Oncology Group. Cancer 2021;127:946–56

4. Arndt CA, Stoner JA, Hawkins DS, Rodeberg DA, Hayes-Jordan AA, Paidas CN, et al. Vincristine, actinomycin, and cyclophosphamide compared with vincristine, actinomycin, and cyclophosphamide alternating with vincristine, topotecan, and cyclophosphamide for intermediate-risk rhabdomyosarcoma: children’s oncology group study D9803. J Clin Oncol 2009;27:5182–8

5. Skapek SX, Ferrari A, Gupta AA, Lupo PJ, Butler E, Shipley J, et al. Rhabdomyosarcoma. Nat Rev Dis Primers 2019;5:1

6. Hsieh J, Danis EP, Owens CR, Parrish JK, Nowling NL, Wolin AR, et al. Dependence of PAX3-FOXO1 chromatin occupancy on ETS1 at important disease-promoting genes exposes new targetable vulnerability in Fusion-Positive Rhabdomyosarcoma. Oncogene 2025;44:19–29

7. Laubscher D, Gryder BE, Sunkel BD, Andresson T, Wachtel M, Das S, et al. BAF complexes drive proliferation and block myogenic differentiation in fusion-positive rhabdomyosarcoma. Nat Commun 2021;12:6924

8. Gryder BE, Yohe ME, Chou HC, Zhang X, Marques J, Wachtel M, et al. PAX3-FOXO1 Establishes Myogenic Super Enhancers and Confers BET Bromodomain Vulnerability. Cancer Discov 2017;7:884–99

9. Wachtel M, Schafer BW. PAX3-FOXO1: Zooming in on an "undruggable" target. Semin Cancer Biol 2018;50:115–23

10. Zhang S, Wang J, Liu Q, McDonald WH, Bomber ML, Layden HM, et al. PAX3-FOXO1 coordinates enhancer architecture, eRNA transcription, and RNA polymerase pause release at select gene targets. Mol Cell 2022;82:4428–42 e7

11. Ahmed AA, Habeebu S, Farooqi MS, Gamis AS, Gonzalez E, Flatt T, et al. MYOD1 as a prognostic indicator in rhabdomyosarcoma. Pediatr Blood Cancer 2021;68:e29085

12. Driman D, Thorner PS, Greenberg ML, Chilton-MacNeill S, Squire J. MYCN gene amplification in rhabdomyosarcoma. Cancer 1994;73:2231–7

13. Pomella S, Cassandri M, D’Archivio L, Porrazzo A, Cossetti C, Phelps D, et al. MYOD-SKP2 axis boosts tumorigenesis in fusion negative rhabdomyosarcoma by preventing differentiation through p57(Kip2) targeting. Nat Commun 2023;14:8373

14. Wang W, Du Y, Datta S, Fowler JF, Sang HT, Albadari N, et al. Targeting the MYCN-MDM2 pathways for cancer therapy: Are they druggable? Genes Dis 2025;12:101156

15. Komori T. Runx2, a multifunctional transcription factor in skeletal development. J Cell Biochem 2002;87:1–8

16. Komori T. Molecular Mechanism of Runx2-Dependent Bone Development. Mol Cells 2020;43:168–75

17. Schroeder TM, Jensen ED, Westendorf JJ. Runx2: a master organizer of gene transcription in developing and maturing osteoblasts. Birth Defects Res C Embryo Today 2005;75:213–25

18. Munehira Y, Yang Z, Gozani O. Systematic Analysis of Known and Candidate Lysine Demethylases in the Regulation of Myoblast Differentiation. J Mol Biol 2017;429:2055–65

19. Pertea M, Kim D, Pertea GM, Leek JT, Salzberg SL. Transcript-level expression analysis of RNA-seq experiments with HISAT, StringTie and Ballgown. Nat Protoc 2016;11:1650–67

20. Bray NL, Pimentel H, Melsted P, Pachter L. Near-optimal probabilistic RNA-seq quantification. Nat Biotechnol 2016;34:525–7

21. Green D, Eyre H, Singh A, Taylor JT, Chu J, Jeys L, et al. Targeting the MAPK7/MMP9 axis for metastasis in primary bone cancer. Oncogene 2020;39:5553–69

22. Love MI, Huber W, Anders S. Moderated estimation of fold change and dispersion for RNA-seq data with DESeq2. Genome Biol 2014;15:550

23. Singh A, Mohorianu I, Green D, Dalmay T, Dasgupta I, Mukherjee SK. Artificially induced phased siRNAs promote virus resistance in transgenic plants. Virology 2019;537:208–15

24. Tattersall L, Shah KM, Lath DL, Singh A, Down JM, De Marchi E, et al. The P2RX7B splice variant modulates osteosarcoma cell behaviour and metastatic properties. J Bone Oncol 2021;31:100398

25. Prufer K, Stenzel U, Dannemann M, Green RE, Lachmann M, Kelso J. PatMaN: rapid alignment of short sequences to large databases. Bioinformatics 2008;24:1530–1

26. Kozomara A, Griffiths-Jones S. miRBase: annotating high confidence microRNAs using deep sequencing data. Nucleic Acids Res 2014;42:D68–73

27. Billmeier M, Green D, Hall AE, Turnbull C, Singh A, Xu P, et al. Mechanistic insights into non-coding Y RNA processing. RNA Biol 2022;19:468–80

28. Ghibaudi M, Boido M, Green D, Signorino E, Berto GE, Pourshayesteh S, et al. miR-7b-3p Exerts a Dual Role After Spinal Cord Injury, by Supporting Plasticity and Neuroprotection at Cortical Level. Front Mol Biosci 2021;8:618869

29. Green D, Mohorianu I, McNamara I, Dalmay T, Fraser WD. miR-16 is highly expressed in Paget’s associated osteosarcoma. Endocr Relat Cancer 2017;24:L27–L31

30. Green D, Singh A, Sanghera J, Jeys L, Sumathi V, Dalmay T, et al. Maternally expressed, paternally imprinted, embryonic non-coding RNA are expressed in osteosarcoma, Ewing sarcoma and spindle cell sarcoma. Pathology 2019;51:113–6

31. Green D, Singh A, Tippett VL, Tattersall L, Shah KM, Siachisumo C, et al. YBX1-interacting small RNAs and RUNX2 can be blocked in primary bone cancer using CADD522. J Bone Oncol 2023;39:100474

32. Shaw B, Burrell CL, Green D, Navarro-Martinez A, Scott D, Daroszewska A, et al. Molecular insights into an ancient form of Paget’s disease of bone. Proc Natl Acad Sci U S A 2019;116:10463–72

33. Li JJ, Kovach AR, DeMonia M, Slemmons KK, Oristian KM, Chen C, et al. Expression of oncogenic HRAS in human Rh28 and RMS-YM rhabdomyosarcoma cells leads to oncogene-induced senescence. Sci Rep 2021;11:16505

34. Ritchie ME, Phipson B, Wu D, Hu Y, Law CW, Shi W, et al. limma powers differential expression analyses for RNA-sequencing and microarray studies. Nucleic Acids Res 2015;43:e47

35. Hanahan D. Hallmarks of Cancer: New Dimensions. Cancer Discov 2022;12:31–46

36. Tanzarella S, Lionello I, Valentinis B, Russo V, Lollini PL, Traversari C. Rhabdomyosarcomas are potential target of MAGE-specific immunotherapies. Cancer Immunol Immunother 2004;53:519–24

37. Hwang GH, Pazyra-Murphy MF, Seo HS, Dhe-Paganon S, Stopka SA, DiPiazza M, et al. A Benzarone Derivative Inhibits EYA to Suppress Tumor Growth in SHH Medulloblastoma. Cancer Res 2024;84:872–86

38. Kim MS, Gernapudi R, Cedeno YC, Polster BM, Martinez R, Shapiro P, et al. Targeting breast cancer metabolism with a novel inhibitor of mitochondrial ATP synthesis. Oncotarget 2020;11:3863–85

39. Kim MS, Gernapudi R, Choi EY, Lapidus RG, Passaniti A. Characterization of CADD522, a small molecule that inhibits RUNX2-DNA binding and exhibits antitumor activity. Oncotarget 2017;8:70916–40

40. Hsu JY, Danis EP, Nance S, O’Brien JH, Gustafson AL, Wessells VM, et al. SIX1 reprograms myogenic transcription factors to maintain the rhabdomyosarcoma undifferentiated state. Cell Rep 2022;38:110323

41. Brusgard JL, Choe M, Chumsri S, Renoud K, MacKerell AD, Jr., Sudol M, et al. RUNX2 and TAZ-dependent signaling pathways regulate soluble E-Cadherin levels and tumorsphere formation in breast cancer cells. Oncotarget 2015;6:28132–50

42. Martin JW, Zielenska M, Stein GS, van Wijnen AJ, Squire JA. The Role of RUNX2 in Osteosarcoma Oncogenesis. Sarcoma 2011;2011:282745

43. Depmap B. DepMap23Q4 Public. Figshare+.Dataset. ed2023.

44. Wei Y, Qin Q, Yan C, Hayes MN, Garcia SP, Xi H, et al. Single-cell analysis and functional characterization uncover the stem cell hierarchies and developmental origins of rhabdomyosarcoma. Nat Cancer 2022;3:961–75

45. Crose LE, Galindo KA, Kephart JG, Chen C, Fitamant J, Bardeesy N, et al. Alveolar rhabdomyosarcoma-associated PAX3-FOXO1 promotes tumorigenesis via Hippo pathway suppression. J Clin Invest 2014;124:285–96

46. Liao GB, Li XZ, Zeng S, Liu C, Yang SM, Yang L, et al. Regulation of the master regulator FOXM1 in cancer. Cell Commun Signal 2018;16:57

47. Kuda M, Kohashi K, Yamada Y, Maekawa A, Kinoshita Y, Nakatsura T, et al. FOXM1 expression in rhabdomyosarcoma: a novel prognostic factor and therapeutic target. Tumour Biol 2016;37:5213–23

48. Bull EC, Singh A, Harden AM, Soanes K, Habash H, Toracchio L, et al. Targeting metastasis in paediatric bone sarcomas. Mol Cancer 2025;24:153

49. Zhang Y, Xie RL, Croce CM, Stein JL, Lian JB, van Wijnen AJ, et al. A program of microRNAs controls osteogenic lineage progression by targeting transcription factor Runx2. Proc Natl Acad Sci U S A 2011;108:9863–8

50. Ebauer M, Wachtel M, Niggli FK, Schafer BW. Comparative expression profiling identifies an in vivo target gene signature with TFAP2B as a mediator of the survival function of PAX3/FKHR. Oncogene 2007;26:7267–81

51. Nomura K, Kimira Y, Osawa Y, Kataoka-Matsushita A, Takao K, Sugita Y, et al. Stimulation of the Runx2 P1 promoter by collagen-derived dipeptide prolyl-hydroxyproline bound to Foxg1 and Foxo1 in osteoblasts. Biosci Rep 2021;41

52. Fritz AJ, Hong D, Boyd J, Kost J, Finstaad KH, Fitzgerald MP, et al. RUNX1 and RUNX2 transcription factors function in opposing roles to regulate breast cancer stem cells. J Cell Physiol 2020;235:7261–72

53. Tuo Z, Zhang Y, Wang X, Dai S, Liu K, Xia D, et al. RUNX1 is a promising prognostic biomarker and related to immune infiltrates of cancer-associated fibroblasts in human cancers. BMC Cancer 2022;22:523

54. Sood R, Kamikubo Y, Liu P. Role of RUNX1 in hematological malignancies. Blood 2017;129:2070–82

55. Akech J, Wixted JJ, Bedard K, van der Deen M, Hussain S, Guise TA, et al. Runx2 association with progression of prostate cancer in patients: mechanisms mediating bone osteolysis and osteoblastic metastatic lesions. Oncogene 2010;29:811–21

56. Khan AS, Campbell KJ, Cameron ER, Blyth K. The RUNX/CBFbeta Complex in Breast Cancer: A Conundrum of Context. Cells 2023;12

57. Kubota S, Tokunaga K, Umezu T, Yokomizo-Nakano T, Sun Y, Oshima M, et al. Lineage-specific RUNX2 super-enhancer activates MYC and promotes the development of blastic plasmacytoid dendritic cell neoplasm. Nat Commun 2019;10:1653

58. Lin TC. RUNX2 and Cancer. Int J Mol Sci 2023;24

59. Sustic T, Bosdriesz E, van Wageningen S, Wessels LFA, Bernards R. RUNX2/CBFB modulates the response to MEK inhibitors through activation of receptor tyrosine kinases in KRAS-mutant colorectal cancer. Transl Oncol 2020;13:201–11

60. Trust BCR. 2023 Revolutionary new bone cancer drug identified. Accessed 2025.

61. Alegre F, Ormonde AR, Godinez DR, Illendula A, Bushweller JH, Wittenburg LA. The interaction between RUNX2 and core binding factor beta as a potential therapeutic target in canine osteosarcoma. Vet Comp Oncol 2020;18:52–63

62. Gayatri MB, Kancha RK, Behera A, Patchva D, Velugonda N, Gundeti S, et al. AMPK-induced novel phosphorylation of RUNX1 inhibits STAT3 activation and overcome imatinib resistance in chronic myelogenous leukemia (CML) subjects. Cell Death Discov 2023;9:401

63. Institute N. 2024 Clinical trial researching therapy for RUNX1 mutations. Accessed 2025.

64. Tuddenham L, Wheeler G, Ntounia-Fousara S, Waters J, Hajihosseini MK, Clark I, et al. The cartilage specific microRNA-140 targets histone deacetylase 4 in mouse cells. FEBS Lett 2006;580:4214–7

65. Greither T, Koser F, Holzhausen HJ, Guttler A, Wurl P, Kappler M, et al. MiR-155-5p and MiR-203a-3p Are Prognostic Factors in Soft Tissue Sarcoma. Cancers (Basel) 2020;12

